# Adaptor interactions trigger Pan1 self-assembly during clathrin-mediated endocytosis in budding yeast

**DOI:** 10.64898/2026.09.04.749396

**Authors:** Mamta Mamta, Anne-Sophie Rivier Cordey, Lazar Ivanovic, V. T. Bagyashree, Wanda Kukulski, Marko Kaksonen

## Abstract

Clathrin-mediated endocytosis requires the coordinated assembly of a highly dynamic protein network that couples membrane remodeling to actin-driven force generation. The essential budding yeast protein, Pan1, scaffolds the endocytic protein network by linking adaptor proteins, and other coat components, and actin assembly regulators. How Pan1 coordinates all these different proteins in space and time is not well understood. We show that Pan1 undergoes biomolecular condensation. Elevated Pan1 expression induced the formation of condensates that exhibited partial rapid molecular exchange, temperature-dependent reversibility, and sensitivity to disruption of weak hydrophobic interactions. The Pan1 assemblies are compositionally selective, preferentially enriching late-stage endocytic factors while excluding early adaptor proteins. Truncation analysis demonstrated that condensation depends on cooperative contributions from intrinsically disordered regions, the EH2 domain, and the oligomerization module, whereas a C-terminal region negatively regulates condensation. Deletion of Pan1 regions promoting its self-assembly perturbed Pan1 assembly and function at endocytic sites. Disruption of interactions between Pan1 and endocytic adaptors did not alter the dynamics of endocytic events but instead reduced the number of endocytic events marked by Pan1 and caused the formation of ectopic Pan1 condensates. This indicates that adaptor-mediated interactions spatially constrain Pan1 localization by seeding Pan1 assembly at endocytic sites. Our findings support a model in which adaptor proteins seed Pan1 assembly at the endocytic sites while multivalent interactions drive Pan1 self-assembly to build the higher-order molecular network of the late endocytic coat.

## Introduction

Clathrin-mediated endocytosis is a conserved membrane trafficking pathway that enables uptake of extracellular cargo, plasma membrane proteins, and lipids while maintaining membrane homeostasis. In *Saccharomyces cerevisiae*, clathrin-mediated endocytosis proceeds through the highly ordered assembly of an extensive, dynamic protein network at cortical sites. Sequential recruitment of coat proteins, adaptors, actin regulators, and membrane-remodeling factors drives membrane invagination and vesicle formation (Goode et al., 2014; Kaksonen et al., 2005; Kaksonen & Roux, 2018; Newpher et al., 2005; Weinberg & Drubin, 2012). Robust execution of this process requires precise spatial and temporal coordination among protein assembly, actin polymerization, and membrane deformation. How these molecular interactions integrate into a coherent, dynamic endocytic structure remains a central question in the field.

A key organizer of the endocytic machinery is Pan1, a scaffold protein and homologue of mammalian intersectin (Tang et al., 1997; Wendland et al., 1998). Deletion of Pan1 is lethal and loss of Pan1 function causes severe defects in cortical actin patch organization, endocytosis, and cell morphogenesis (Bradford et al., 2015). Pan1 physically and functionally associates with End3 and Sla1 to form a complex that regulates cargo capture, coat maturation, and actin assembly (Sun et al., 2015; Tang et al., 2000). In addition, Pan1-End3 complex plays a critical role in coupling Arp2/3- dependent actin assembly to sites of clathrin-mediated endocytosis, thereby linking endocytic coat maturation to force-generating actin network required for membrane invagination (Duncan et al., 2001; Sun et al., 2015, 2017).

Pan1 has two EH domains that bind NPF motif-containing proteins, which include clathrin adaptors Ent1/2 (epsins) and Yap1801/2 (AP180/CALM) (Wendland et al., 1999; Whitworth et al., 2014). Deletion of the adaptor proteins while retaining only their ENTH domain which is required for viability resulted in longer Pan1 lifetimes and impairs endocytic internalization (Claudio Aguilar et al., 2003; Duncan et al., 2001; Maldonado-Báez et al., 2008). Pan1 contains multiple functional domains that mediate interactions with distinct endocytic proteins and previous studies demonstrated that no single domain is sufficient to support Pan1 function, highlighting the importance of coordinated interactions across the protein. Distinct Pan1 regions can interact with one another, raising the possibility that higher-order Pan1 assembly is mediated by distributed intramolecular and/or intermolecular interactions (Miliaras et al., 2004). However, the functional significance of these interactions has remained unclear.

Increasing evidence suggests that biomolecular condensation can contribute to the organization of complex cellular assemblies. Through multivalent interactions, proteins and nucleic acids can assemble into dynamic, non-membrane-bound compartments that selectively concentrate specific factors while excluding others (Banani et al., 2017; Shin & Brangwynne, 2017). Such assemblies have been implicated in diverse cellular processes, including transcriptional regulation, signal transduction, and cytoskeletal organization. However, the extent to which liquid-liquid phase separation underlies the formation and function of cellular assemblies remains actively debated. Commonly reported features, including spherical morphology, fusion, fluorescence recovery after photobleaching, and sensitivity to 1,6-hexanediol, are not individually sufficient to establish phase separation, as similar behaviors may arise from multivalent binding, polymerization, gelation, or other forms of higher-order self-assembly. Consequently, rigorous interpretation requires complementary experimental evidence and careful distinction between phase separation and the broader phenomenon of biomolecular condensation (Alberti et al., 2019; McSwiggen et al., 2019; Musacchio, 2022).

Recent work has implicated condensation in endocytosis. Endocytic proteins containing prion-like domains can form viscoelastic assemblies that contribute directly to membrane remodeling, suggesting that higher-order protein organization may perform mechanical functions beyond simply concentrating molecules (Bergeron-Sandoval et al., 2021). Similarly, the mammalian early endocytic scaffold Eps15 and its budding yeast homolog Ede1 form condensates, and perturbation of their multivalent interactions affects endocytic-site initiation and maturation (Boeke et al., 2014; Day et al., 2021; Kozak & Kaksonen, 2022). Together, these observations suggest that condensation or related forms of multivalent self-assembly may represent broader organizing principles within the endocytic pathway.

It is unknown whether Pan1’s extensive interaction network reflects a simple collection of pairwise binding events or instead arises through higher-order self-assembly mechanisms that promote selective, spatially restricted organization of endocytic factors. Moreover, the relationship between canonical adaptor-mediated recruitment pathways and potential condensate behavior has not been explored.

## Results

### Pan1 forms dynamic concentration-dependent condensates in cells

Recent studies suggest that biomolecular condensation contributes to the organization of the endocytic machinery, specifically the orthologous early-arriving proteins, Ede1 and Eps15. We therefore asked whether Pan1 might have a similar biomolecular property of condensation because the two proteins share several sequence and architectural features. Both are large multidomain scaffolds containing N-terminal EH domains, extensive intrinsically disordered and low-complexity regions, numerous short interaction motifs, and oligomerization elements (Fig. S1A).

We first examined the sequence properties of Pan1 using multiple independent predictors of disorder, interaction potential, and condensation propensity. Sequence-based analyses consistently indicated a strong intrinsic propensity of Pan1 for higher-order assembly (Fig. 1A). Pan1 ranked highly for prion-like sequence composition (PLAAC), intrinsic disorder (ESpritz-DisProt), low-complexity regions (SEG), granule-forming propensity (catGRANULE), π-π interaction potential (PScore), and both droplet- (PdPS) and aggregation-promoting (SaPS) features, suggesting that Pan1 may possess the capacity to access different higher-order assembly states rather than being restricted to a single type of molecular organization. In addition, Pan1 exhibited low hydropathy and a low proportion of charged residues, characteristics commonly associated with intrinsically disordered proteins that undergo biomolecular condensation. Consistent with these sequence features, PhasePred predicted a high propensity for spontaneous self-assembly (PS-Self score = 0.959).

**Figure 1.**
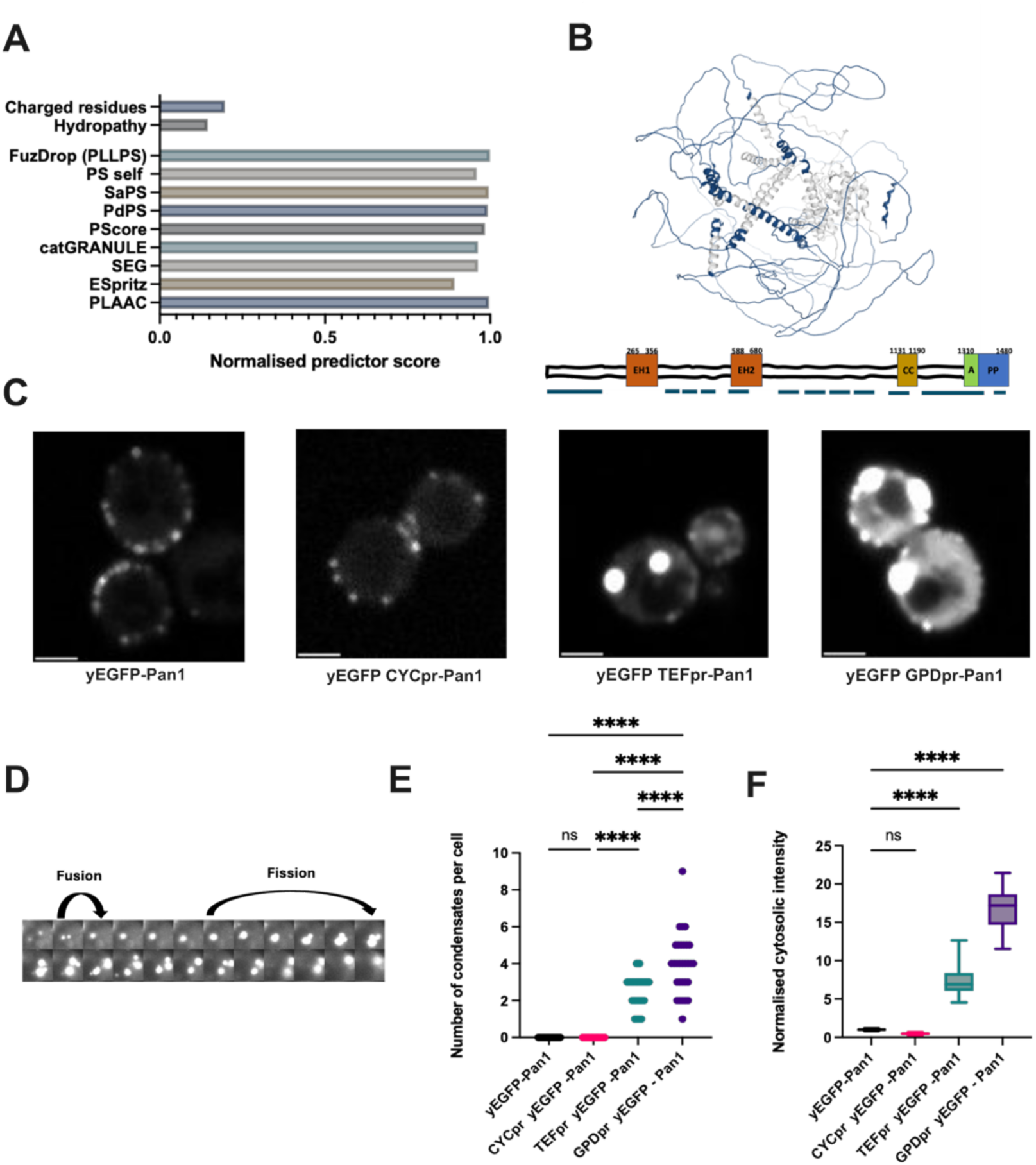
**A**) Horizontal bar plot showing normalized sequence-feature and condensation-prediction scores for Pan1. **(B)** Predicted structural model and domain organization of Pan1. The domain schematic highlights the structured regions of Pan1, including the two N-terminal Eps15 homology domains (EH1 and EH2), the coiled-coil (CC) domain and the C-terminal regulatory region. The remaining regions of the protein are predicted to be predominantly intrinsically disordered (drawn in black). Blue dashed outlines indicate droplet-promoting regions predicted by FuzDrop, corresponding to the blue-highlighted segments in the structural model. **(C)** Representative fluorescence images of GFP tagged Pan1 expressed under promoters of increasing strength. Scale bar, 5 µm. **(D)** Time lapse images showing dynamic fusion and fission behavior of Pan1 condensates over 24 s. **(E)** Quantification of condensate number per cell under increasing promoter strengths. Statistical significance was determined by one-way ANOVA, ****P < 0.0001, N = 30. **(F)** Quantification of normalized cytoplasmic Pan1 intensity under increasing promoter strengths. Statistical significance was determined by one-way ANOVA, ****P < 0.0001, N = 30.

Independent analysis using FuzDrop further predicted an exceptionally high probability of spontaneous liquid-liquid phase separation (PLLPS = 0.9999) and supports the notion that Pan1 possesses an intrinsic capacity for higher-order self-assembly. To identify which regions, underlie this propensity, FuzDrop analysis identified multiple discrete condensation-promoting segments highlighted in teal blue and represented by a blue dashed line in the representative domain organization (Fig. 1B). These regions are distributed throughout the Pan1 sequence and largely overlap with intrinsically disordered regions, suggesting that condensation propensity is encoded in a distributed manner rather than within a single defined domain.

Consistent with this, the AlphaFold2-predicted structural model, together with sequence-based disorder predictions, showed that Pan1 consists of extensive intrinsically disordered regions interspersed with a limited number of folded domains, i.e. the tandem EH domains and the coiled-coil region (Fig. 1B). This architecture, combining discrete folded interaction modules with large, disordered segments, is characteristic of proteins that undergo multivalent self-assembly and biomolecular condensation (Alberti et al., 2019; Banani et al., 2017). These computational analyses suggest that Pan1 possesses a propensity for higher-order self-assembly.

These predictions prompted us to test whether Pan1 exhibits condensation in cells. We replaced the de novo Pan1 promoter with artificial promoters of increasing strength (CYCpr < TEFpr < GPDpr) (Ralser et al., 2007).When Pan1-yEGFP was expressed either under the wild-type promoter or moderately overexpressed with the CYC1 promoter it was localized to endocytic sites at the plasma membrane. Endogenous Pan1 patches appear as small, diffraction-limited cortical puncta that assemble transiently at the plasma membrane and disappear within tens of seconds as endocytosis progresses. Further increases in expression under the stronger TEF1 and GPD promoters led to a pronounced accumulation of Pan1 into distinct assemblies throughout the cell. Because the signal intensity of these assemblies greatly exceeded that of endocytic puncta, imaging conditions were adjusted to partially saturate the assembly signal to allow simultaneous visualization of the comparatively dim endocytic sites within the same image (Figure 1C). Unlike endocytic sites, which are consistently associated with the plasma membrane, these assemblies were frequently observed in the cytoplasm and were not restricted to membrane-proximal regions. The Pan1 assemblies were observed exclusively when Pan1 expression was driven by a strong heterologous promoter and were not detected at endogenous expression levels. We therefore interpret these assemblies as an experimentally induced phenotype reflecting the propensity of Pan1 to assemble rather than enlarged versions of physiological endocytic patches. Given that PAN1 is essential for viability, precluding conventional loss-of-function approaches, we leveraged this overexpression-induced phenotype as an alternative strategy to probe its function. Although these assemblies likely represent nonphysiological structures, their robust and reproducible formation suggests that they arise from intrinsic biochemical properties of Pan1, such as multivalency or an inherent propensity for phase separation.

We next asked whether assembly formation was a general feature of increasing endocytic protein concentration, or whether Pan1 and Ede1 showed an unusual response to elevated expression. To address this, we examined a representative set of proteins from different modules of the endocytic pathway under overexpression conditions (Abp1, Las17, Myo5, and Pan1). At endogenous expression levels, the fluorescently tagged proteins localized to discrete cortical puncta, consistent with their established localization at sites of clathrin-mediated endocytosis. Upon overexpression, the proteins examined retained normal punctate localization, accompanied by increased diffuse cytoplasmic fluorescence. However, none of the other endocytic proteins tested formed large assemblies comparable to those produced by Pan1 (Fig. S1B).

To further characterize Pan1 assemblies, we performed time-lapse live-cell imaging. The assemblies were long lived and over the course of imaging, they exhibited dynamic behaviors, including apparent fusion of neighboring assemblies into larger structures and fission events that generated smaller assemblies (Figure 1D, Movie S1). These observations indicate that Pan1 assemblies are highly dynamic and can continuously reorganize. The number of Pan1 assemblies per cell increased with promoter strength, indicating a concentration dependence. (Figure 1E). To test whether the cytosolic concentration of Pan1 is buffered to a critical level, we quantified the cytosolic intensity of fluorescently tagged Pan1 expressed with different promoters. The cytosolic intensity of Pan1 increased with increasing promoter strength (Figure 1F). This indicates that the Pan1 assembly does not exhibit a dilute-phase concentration buffering expected for a simple phase separation process. Because fluorescent protein tags can potentially promote protein clustering or alter the assembly behavior of intrinsically disordered proteins, we sought to exclude the possibility that Pan1 assembly formation resulted from a tagging artifact. To address this, we examined Pan1 localization using alternative fluorophores and tag orientations. Pan1 tagged with mNeonGreen and expressed from the TEF1 promoter (TEF1pr-mNG-Pan1) formed assemblies comparable to those seen with yEGFP, indicating that assembly formation is not specific to the yEGFP tag (Fig. S1C). Similarly, a C-terminally tagged construct (Pan1-yEGFP) overexpressed from the TEF1 promoter also formed assemblies, indicating that the phenomenon is independent of tag position (Fig. S1D).

### Pan1 forms dynamic and reversible condensates

To determine whether Pan1 assemblies exhibit properties characteristic of biomolecular condensates, we first examined their dynamic behavior using fluorescence recovery after photobleaching (FRAP). Pan1 assemblies recovered rapidly after photobleaching with a half-time of 1.5 s and a mobile fraction of ∼40% (Fig. 2A), indicating substantial molecular exchange between the assemblies and the surrounding cytoplasm. This partial rapid recovery indicates that Pan1 assemblies behave as dynamic condensates containing both mobile and relatively immobile molecular populations.

**Figure 2.**
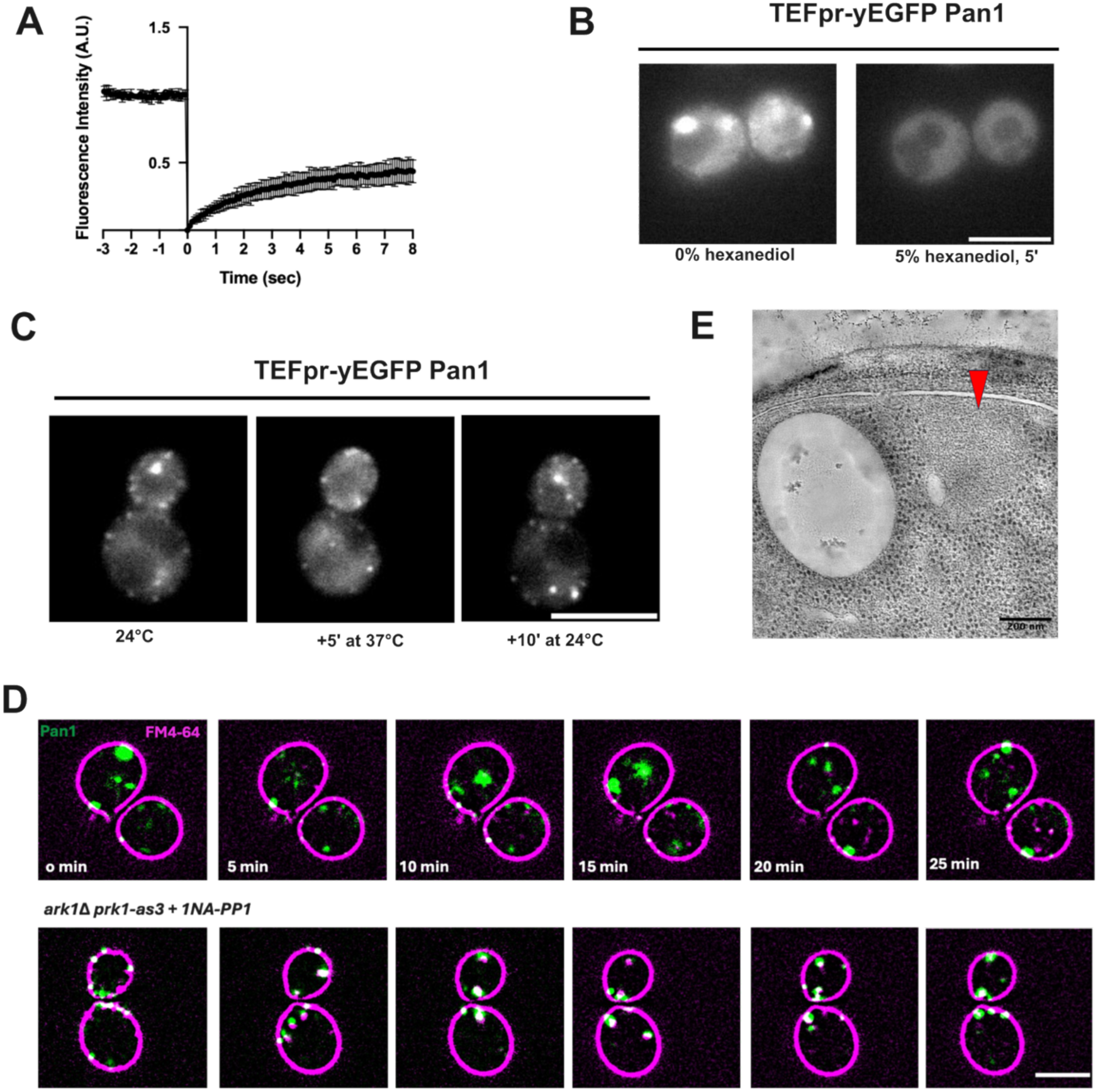
**(A**) Fluorescence recovery after photobleaching (FRAP) curve of TEFpr -yEGFP-Pan1 condensates. **(B)** Representative fluorescence images of TEFpr-yEGFP-Pan1 cells before and after treatment with 5% 1,6- hexanediol. Scale bar, 5 µm. **(C)** Temperature-dependent reversibility of Pan1 condensates. Condensates disappear following incubation at 37°C and rapidly re-form upon return to 24°C. Scale bar, 5 µm. **(D)** Localization of FM4-64 relative to Pan1 condensates. Upper panel: FM4-64 does not appreciably accumulate within Pan1 condensates formed upon Pan1 overexpression. Lower panel: FM4-64 accumulates in Pan1-containing aggregates in *ark1Δ prk1-as3* cells following kinase inhibition. Scale bar, 5 µm. **(E)** Correlative light and electron microscopy (CLEM) analysis of Pan1 condensates. Condensates correspond to electron-dense ribosome-excluding regions lacking detectable membrane structures (region pointed by red arrow).

We next tested the sensitivity of Pan1 assemblies to 1,6-hexanediol which is an aliphatic alcohol that is commonly used to perturb weak interactions associated with some condensates. Treatment with 5% 1,6-hexanediol resulted in rapid dissolution of Pan1 assembly within five minutes (Fig. 2B), indicating that condensate integrity depends on weak multivalent interactions.

Because condensate formation is often influenced by environmental conditions, we examined the effect of temperature on Pan1 assembly. Increasing the temperature to 37°C led to rapid disappearance of condensates, whereas returning cells to 24°C restored condensate formation within ten minutes (Fig. 2C), indicating that Pan1 assembly is reversible and temperature dependent.

We test whether Pan1 condensates resemble previously reported Pan1-containing aggregates observed in *ark1Δ prk1-as3* mutants (Sekiya-Kawasaki et al., 2003). Ark1 and Prk1 are endocytic kinases that phosphorylate Pan1 and other coat components to promote disassembly of the endocytic machinery following vesicle internalization. Loss of Ark1/Prk1 activity prevents efficient coat disassembly, resulting in the accumulation of large membrane-associated Pan1 containing aggregates enriched in endocytic proteins and internalized membrane. These structures can be readily visualized by the lipophilic dye FM4-64.

FM4-64 strongly accumulated in Pan1 containing structures in *ark1Δ prk1-as3* cells following kinase inhibition, consistent with previous reports. In contrast, FM4-64 did not appreciably colocalize with Pan1 condensates formed upon Pan1 overexpression (Fig. 2D), indicating that these assemblies are compositionally distinct from the vesicle associated endocytic aggregates.

To further characterize the ultrastructural organization of Pan1 condensates, we performed correlative light and electron microscopy (CLEM). On sections of resin-embedded cells overexpressing Pan1, we acquired electron tomograms in regions containing Pan1-yEGFP signals indicative of condensates. Pan1 condensates corresponded to regions that were largely devoid of ribosomes and lacked detectable membrane structures (Fig. 2E, Movie S2), consistent with the formation of a protein-rich, membrane-free compartment. This ultrastructural organization resembles previously described Ede1 condensates which similarly form ribosome-excluding compartments (Wilfling et al., 2020).

Together, these findings demonstrate that Pan1 assemblies exhibit the features commonly associated with biomolecular condensates including rapid molecular exchange, sensitivity to weak interaction perturbation and environmental responsiveness. Collectively, these observations suggest that Pan1 condensates represent regulated biomolecular assemblies different from static protein aggregates.

### Pan1 condensates selectively recruit a subset of endocytic proteins

Because Pan1 functions as a central scaffold that interacts with proteins from several endocytic modules, we asked whether the Pan1 condensates retain the ability to recruit other components of the endocytic machinery. To characterize the molecular composition of Pan1 condensates, we examined the localization of fluorescently tagged endocytic proteins relative to Pan1 assemblies in cells overexpressing Pan1 (Figure 3A).

**Figure 3.**
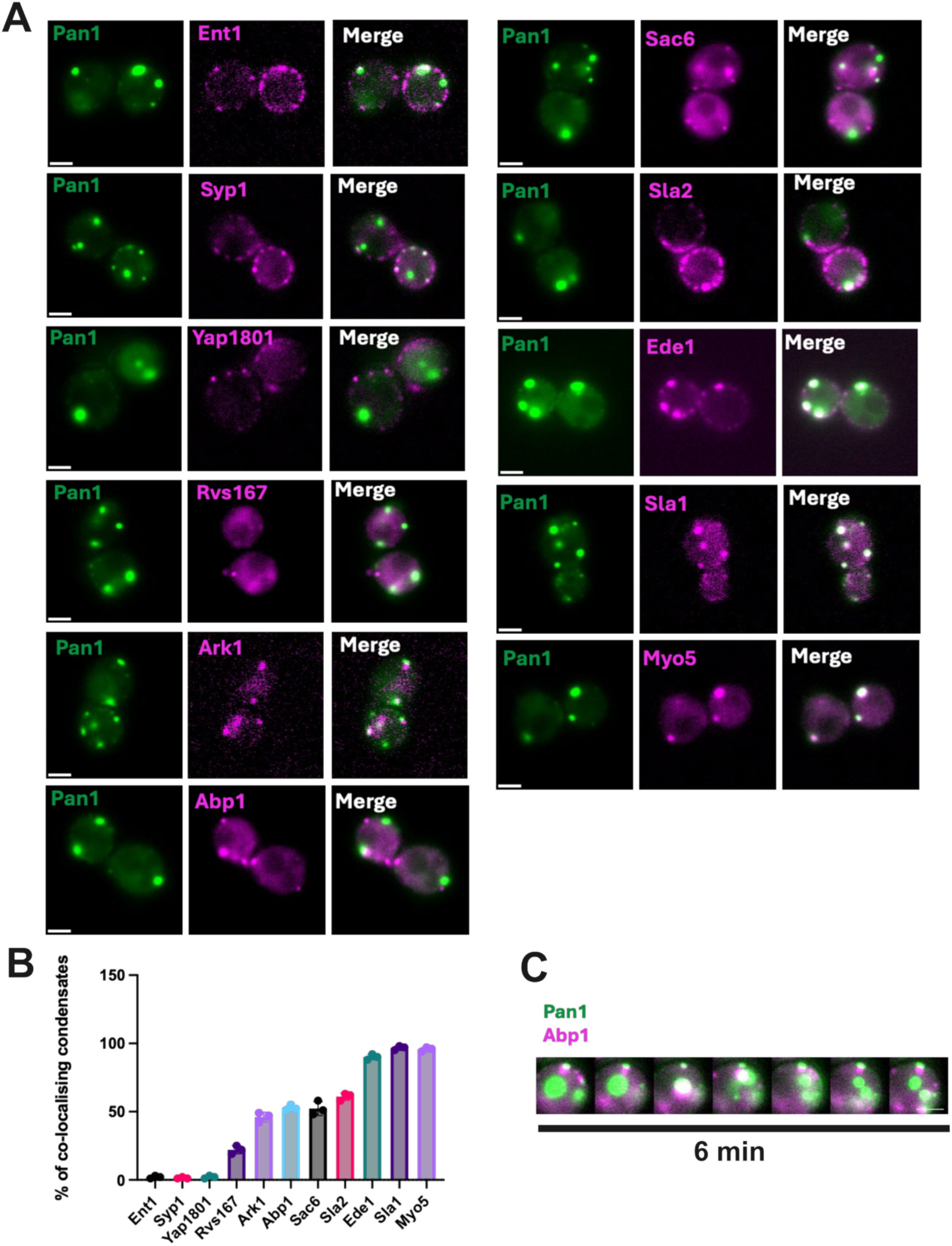
(**A**) Representative fluorescence images showing localization of endocytic proteins relative to Pan1 condensates in cells overexpressing Pan1. Scale bars, 2 µm. **(B)** Quantification of colocalization of endocytic proteins within Pan1 condensates, revealing differential partitioning of endocytic factors. Statistical significance was determined by one-way ANOVA, ****P < 0.0001, N = 30. **(C)** Time-lapse images showing transient recruitment of Abp1 to Pan1 condensates over 6 min. Scale bar, 2 µm.

The early adaptor proteins including the Ent1, Yap1801 and Syp1 (Fcho1/2) showed little or no enrichment within Pan1 condensates. In contrast, several factors associated with later stages of endocytosis were efficiently recruited. The adaptor Sla1 and the type I myosin, Myo5 showed strong enrichment within Pan1 condensates, consistent with their direct interaction with Pan1. The early-arriving scaffold Ede1, which is recruited before Pan1 during endocytosis was also robustly incorporated, indicating that Pan1 assemblies can recruit upstream organizational components of the endocytic machinery.

Quantitative analysis further revealed differential partitioning among late-stage factors (Figure 3B). Sla2 displayed enrichment within condensates, suggesting preferential incorporation of proteins linking the endocytic coat to the actin cytoskeleton. By contrast, the amphiphysin like protein Rvs167, which functions during membrane invagination and scission, exhibited comparatively weaker enrichment. Similarly, the kinase Ark1 showed only limited partitioning into Pan1 condensates. These analyses revealed that Pan1 condensates are compositionally selective, preferentially recruiting a specific subset of proteins while excluding others.

Interestingly, the actin binding protein, Abp1, was only transiently associated with Pan1 condensates (Figure 3C). In many cases, Abp1 recruitment coincided with or immediately preceded condensate fission or dissolution events, suggesting that actin assembly may contribute to remodeling or disassembly of Pan1 condensates.

To further characterize the relationship between Pan1 condensates and the actin cytoskeleton, we examined the localization of filamentous actin using phalloidin staining. Phalloidin-positive structures were occasionally observed adjacent to or partially associated with Pan1 condensates (Figure S2A), consistent with the dynamic association of actin related factors observed for Abp1 (Figure 3C).

Because condensation of Ede1 is proposed to initiate endocytosis, we next asked whether Pan1 condensate formation depends on Ede1. Pan1 condensates remained readily detectable in an *ede1Δ* background following Pan1 overexpression indicating that Ede1 is not required for Pan1 condensation (Figure S2B).

To further exclude fluorescent-tagging artifacts, we analyzed cells expressing overexpressed untagged Pan1 together with Abp1-mCherry. Because Abp1 transiently associates with Pan1 condensates without stable incorporation, the appearance of large Abp1-positive structures provides an indirect readout of Pan1 condensate formation, further supporting the conclusion that Pan1 condensation is independent of fluorescent tagging (Figure S2C).

### Multiple regions of Pan1 cooperatively drive and regulate condensate formation

To identify the regions in the sequence responsible for Pan1 condensation, we performed truncation analysis. Because *PAN1* is essential for viability, precluding conventional loss-of-function approaches, we generated a series of heterozygous truncation mutants in a diploid background in which one allele remained wild type while the second allele encoded the indicated truncation, allowing region-specific interrogation of Pan1 assembly while preserving an essential wild-type copy of the gene.

Consistent with our earlier results, overexpression of full-length Pan1 robustly induced cytoplasmic condensates (Fig. 4A). Deletion of the N-terminal intrinsically disordered region (Δ2-265) eliminated condensate formation (Fig. 4B), and extending this deletion to include both EH domains (Δ2-680) similarly prevented assembly (Fig. 4G), together indicating that this N-terminal region is required for condensation.

**Figure 4.**
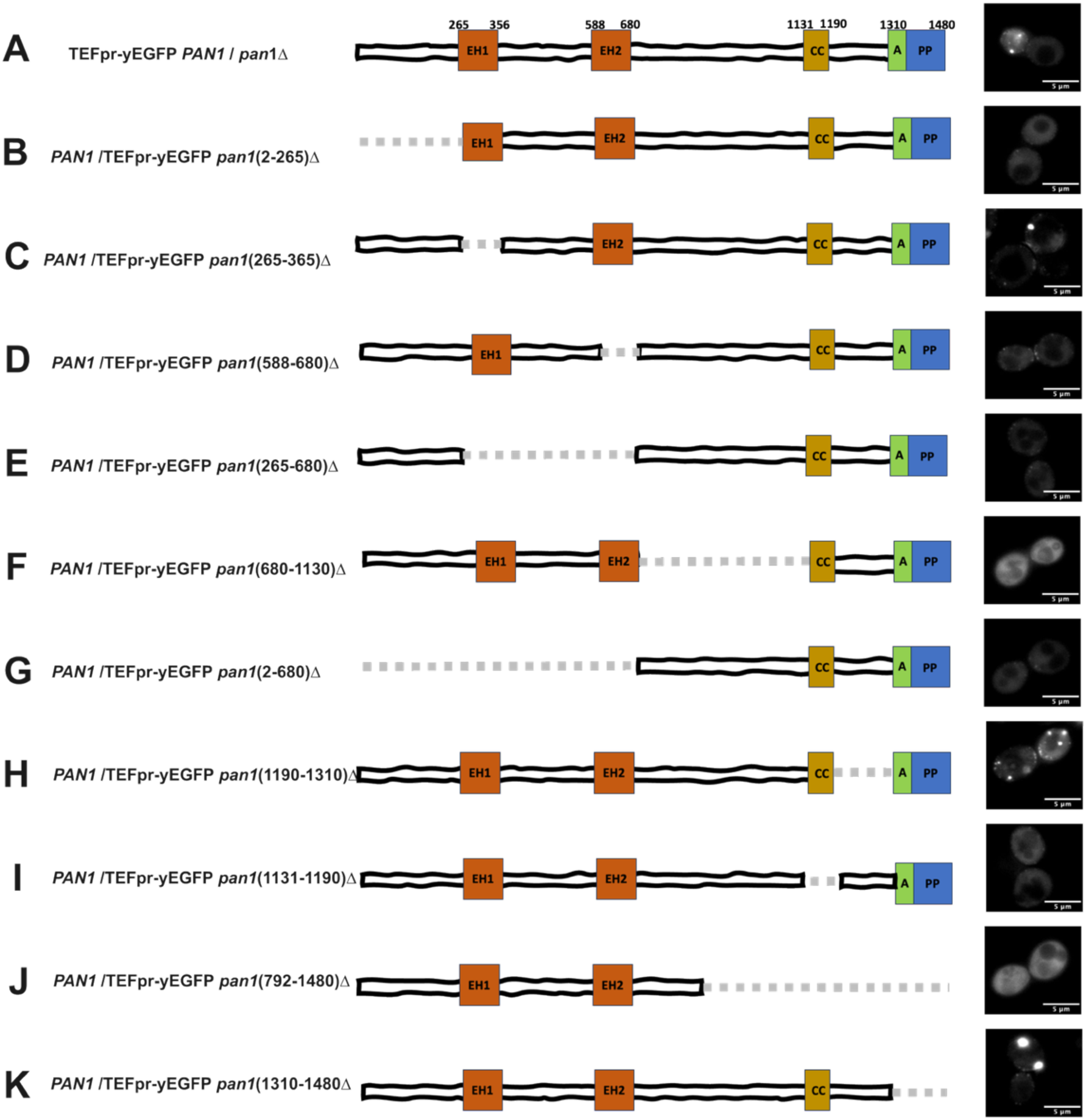
**(A-K)** Schematic representation of Pan1 truncation constructs analyzed in a heterozygous diploid background together with representative fluorescence images showing condensate phenotypes following overexpression of each construct. Deleted regions are indicated by dashed segments. EH1, EH2, coiled-coil (CC), actin-binding (A), and proline-rich (PP) regions are indicated. **(A)** Full-length TEFpr-yEGFP-Pan1 **(B)** Deletion of the N-terminal intrinsically disordered region (Δ2- 265). **(C)** Deletion of EH1 (Δ265-365) **(D)** Deletion of EH2 (Δ588-680) **(E)** Deletion of the EH-containing region (Δ265-680) **(F)** Deletion of the central region (Δ680-1130) **(G)** Deletion of the N-terminal region including both EH domains (Δ2-680) **(H)** Deletion of residues 1190-1310 **(I)** Deletion of the coiled-coil domain (Δ1131-1190) **(J)** Deletion of residues 792-1480 **(K)** Deletion of the C-terminal region (Δ1310-1480)

We next examined the two EH domains individually. Deletion of EH1 (Δ265-365) had no effect on condensate formation (Fig. 4C), whereas deletion of EH2 (Δ588-680) abolished it entirely (Fig. 4D), as did removal of the full EH-containing region (Δ265-680) (Fig. 4E). These results identified EH2, but not EH1, as a critical determinant of condensation, indicating that the two EH domains are functionally distinct in this context. Deletion of the central region (Δ680-1130) also abolished assembly (Fig. 4F), as did the larger truncation Δ792-1480, which spans this region (Fig. 4J), indicating that the central disordered segment makes a critical contribution to multivalent assembly. Deletion of the coiled-coil domain alone (Δ1131-1190) likewise eliminated condensate formation (Fig. 4I), consistent with a role for coiled-coil-mediated oligomerization in driving Pan1 self-assembly.

In contrast to these assembly-promoting elements, we identified a C-terminal region that negatively regulates condensation. Deletion of residues 1190-1310 increased condensate formation (Fig. 4H), and deletion of the larger, non-overlapping C-terminal segment spanning residues 1310-1480 which leaves the coiled-coil domain intact produced condensates that were both larger and more numerous than those formed by full-length Pan1 (Fig. 4K, S3A, S3B). Because this region mediates interactions with SH3-domain containing type I myosins (Barker et al., 2007), these results raise the possibility that myosin binding restrains Pan1 condensation *in vivo*.

Together, these findings demonstrate that Pan1 assembly is governed by a balance of positive and negative regulatory inputs. Whereas the N-terminal intrinsically disordered region, EH2 domain, central region, and coiled-coil domain promote assembly, the C-terminal myosin-interacting region limits condensation. This regulatory architecture suggests that Pan1 assembly is dynamically modulated during endocytosis through coordinated multivalent interactions and inhibitory binding events.

To determine whether differences in protein expression could explain the condensate phenotypes observed for the Pan1 truncation mutants, we compared the abundance of all GFP-tagged constructs by immunoblotting with an anti-GFP antibody, using PGK1 as a loading control (Fig. S3C). Although significant differences in protein abundance were observed between individual constructs, these did not correlate with their ability to form condensates. For example, several mutants that failed to assemble into condensates were expressed at levels comparable to (Fig 4D, E) or higher than (Fig 4J) constructs that readily condensed, whereas some strongly condensing mutants were expressed at relatively moderate levels (Fig 4K) (Fig. S3D). These results indicate that the distinct condensation phenotypes of the Pan1 truncation mutants arise from disruption of specific sequence elements required for higher order assembly.

Previous yeast two-hybrid analysis suggested that distinct regions of Pan1 can interact with one another (Miliaras et al., 2004). To more comprehensively define these interaction networks, particularly within the intrinsically disordered regions implicated in condensation, we performed an expanded yeast two-hybrid analysis including the intrinsically disordered regions (Fig. S4A). Yeast two-hybrid analysis revealed interactions between distinct Pan1 fragments, consistent with multiple weak interaction interfaces distributed throughout the protein (Fig. S4A). Specifically, yeast two-hybrid analysis identified IDR1 (1-263) and IDR2 (359-593) as two of the main regions mediating multiple interactions with other Pan1 regions, suggesting that distributed weak multivalent interactions mediated by these IDRs contribute to higher order Pan1 assembly. Consistent with this, the protein region mapping experiment identified two key intrinsically disordered regions that contribute to Pan1 condensation: the N-terminal IDR1 (2-265) and a second intrinsically disordered region located between the EH domains: IDR2 (365-588).

Together, the requirement for the intrinsically disordered regions, EH2 domain, and coiled-coil domain for condensate formation suggests that Pan1 condensation is driven by distributed multivalent interactions rather than a single dedicated assembly region. Such an organization would enable dynamic and reversible scaffold assembly while remaining highly responsive to regulatory interactions like myosin binding during endocytosis. These multivalent interactions may allow the coat to remain dynamic and rearrangeable during assembly while acquiring greater structural constraint as the endocytic machinery transitions to actin-driven membrane invagination (Boinet et al., 2026).

### IDR-mediated multivalent interactions regulate Pan1 assembly dynamics at endocytic sites

Having established that IDR-mediated multivalent interactions are critical for Pan1 condensation in the overexpression assay, we next asked whether these same interactions regulate assembly of the endogenous late endocytic coat when Pan1 is expressed at endogenous level. To address this, we generated haploid strains lacking either IDR1 (Δ2-265) or IDR2 (Δ365-588) and monitored Pan1 cortical patch dynamics by live-cell imaging.

Deletion of IDR1 increased the mean lifetime of Pan1 cortical patches from 29.5 s in wild-type cells to 41.5 s and broadened the distribution of patch lifetimes (Fig. 5A, 5B). Deletion of IDR2 produced a more pronounced phenotype, increasing the mean lifetime to 62.6 s and further increasing the variability of Pan1 dynamics. Consistent with these observations, the coefficient of variation increased progressively from 18.8% in wild-type cells to 31.1% following IDR1 deletion and to approximately 42% following IDR2 deletion, indicating that loss of IDR mediated interactions compromises the efficiency and regularity of Pan1 assembly at endocytic sites with IDR2 playing a major role compared to IDR1.

**Figure 5.**
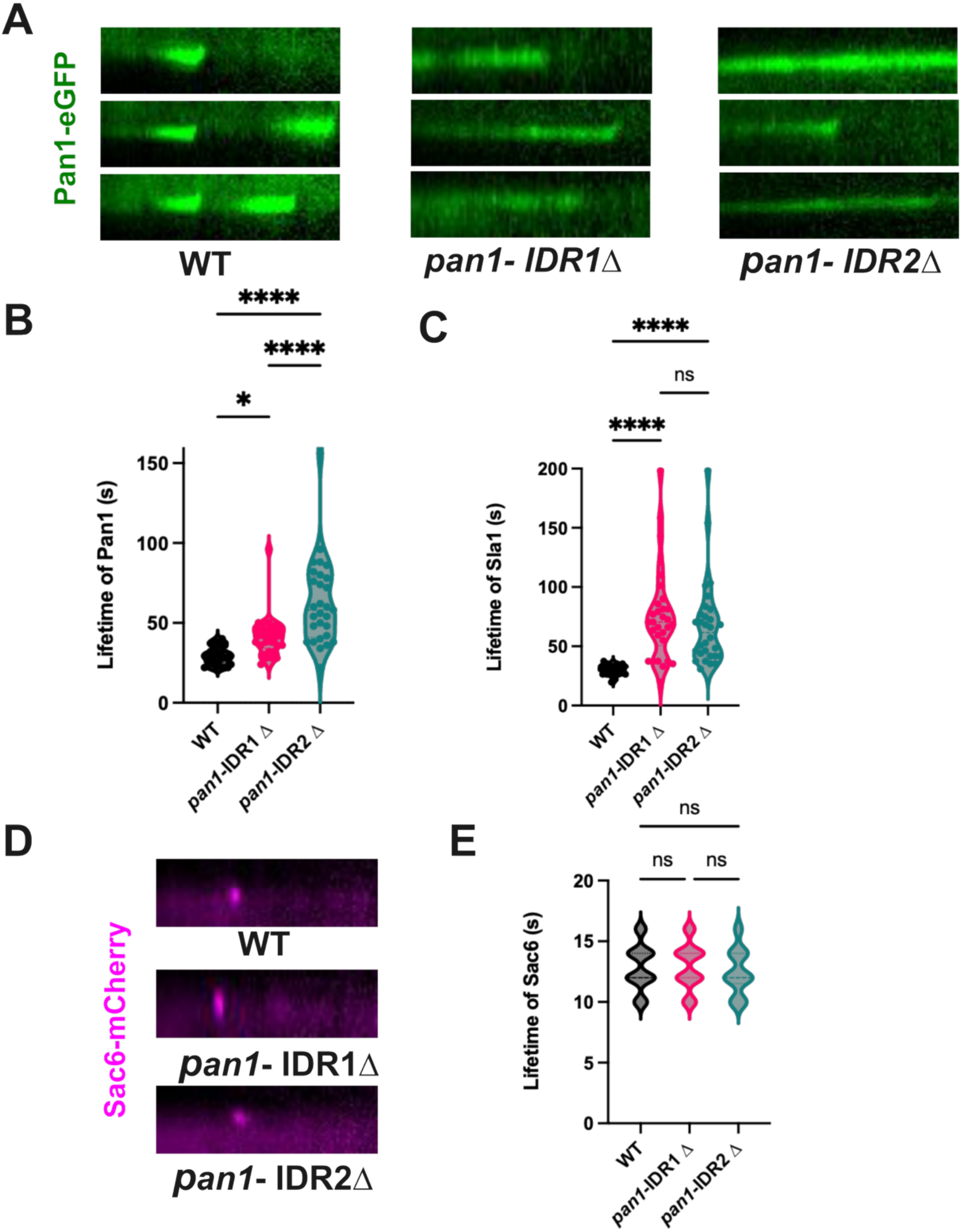
(**A**) Representative kymographs of Pan1-eGFP cortical patches in wild-type (WT), *pan1*-ΔIDR1 (Δ2-265), and *pan1*-ΔIDR2 (Δ365-588) cells. **(B)** Quantification of Pan1 cortical patch lifetimes in WT, *pan1*-ΔIDR1, and *pan1*-ΔIDR2 cells. Violin plots show the distribution of individual patch lifetimes (n = 30 endocytic events). Brown-Forsythe ANOVA, P < 0.0001**. (C)** Quantification of Sla1 cortical patch lifetimes in WT and the indicated IDR deletion mutants. Violin plots show the distribution of individual patch lifetimes (n = 30 endocytic events). Brown-Forsythe ANOVA, P = 0.0002**. (D)** Representative kymographs of Sac6-mCherry cortical patches in WT, Pan1-ΔIDR1, and Pan1-ΔIDR2 cells. **(E)** Quantification of Sac6 cortical patch lifetimes in WT and the indicated IDR deletion mutants. Violin plots show the distribution of individual patch lifetimes (n=30 endocytic events) One-way ANOVA, P = 0.5815 (not significant).

We next sought whether these defects were propagated through the late endocytic coat. Both IDR1 and IDR2 deletions significantly prolonged the lifetime of the late coat protein Sla1, with comparable effects observed for the two mutants (Fig. 5C). These findings indicate that disruption of Pan1- mediated multivalent interactions affects the dynamics of the late coat network rather than being restricted to Pan1 itself.

Strikingly, the lifetime of the late actin marker Sac6 remained indistinguishable from wild type in both mutants (Fig. 5D, 5E). Thus, disruption of IDR-mediated interactions selectively delays assembly and maturation of the late coat without affecting the kinetics of the downstream actin module. A possible explanation is that loss of these weak multivalent interactions slows accumulation of a functional Pan1 scaffold, thereby prolonging the residence of late coat components. Once sufficient Pan1 has assembled, however, recruitment and turnover of the actin machinery proceed with normal kinetics.

These findings demonstrate that the same intrinsically disordered regions required for Pan1 condensation upon overexpression also regulate the dynamics of the endogenous late endocytic scaffold. Thus, IDR-mediated multivalent interactions are not merely revealed by ectopic overexpression but instead promote efficient, robust, and well-timed assembly of the late endocytic coat during clathrin-mediated endocytosis.

### EH-NPF interactions trigger Pan1 assembly without affecting endocytic kinetics

Our findings so far establish that Pan1 can undergo higher order self-assembly driven by weak multivalent interactions, and that these same interactions contribute to robust assembly of the endogenous late endocytic coat. This raised the question of how Pan1 assembly is spatially restricted to endocytic sites rather than occurring throughout the cytoplasm. A plausible mechanism is Pan1 recruitment through endocytic adaptor proteins. The epsins Ent1/2 and the AP180 homologs Yap1801/2 bind the N-terminal EH domains of Pan1 through multiple NPF motifs, making these interactions attractive candidates for locally triggering Pan1 assembly at sites of clathrin-mediated endocytosis.

To examine the contribution of the EH-NPF interactions on the recruitment and endocytic dynamics of Pan1, we generated a series of strains in which increasing numbers of NPF motifs were mutated to NAA across different combinations of Pan1 binding adaptors Ent1, Ent2, Yap1801, and Yap1802. These included a mutant in which all four NPF motifs present across Ent1 and Ent2 were substituted with NAA (4NPF-NAA), a mutant carrying all ten NPF-to-NAA substitutions in Yap1801 and Yap1802 (10NPF-NAA), a combined Ent1/Ent2/Yap1801 mutant containing nine NPF-to-NAA substitutions (9NPF-NAA), and a strain in which all fourteen NPF motifs across the Ent and Yap adaptor families were mutated (14NPF-NAA) (Fig. 6A).

**Figure 6.**
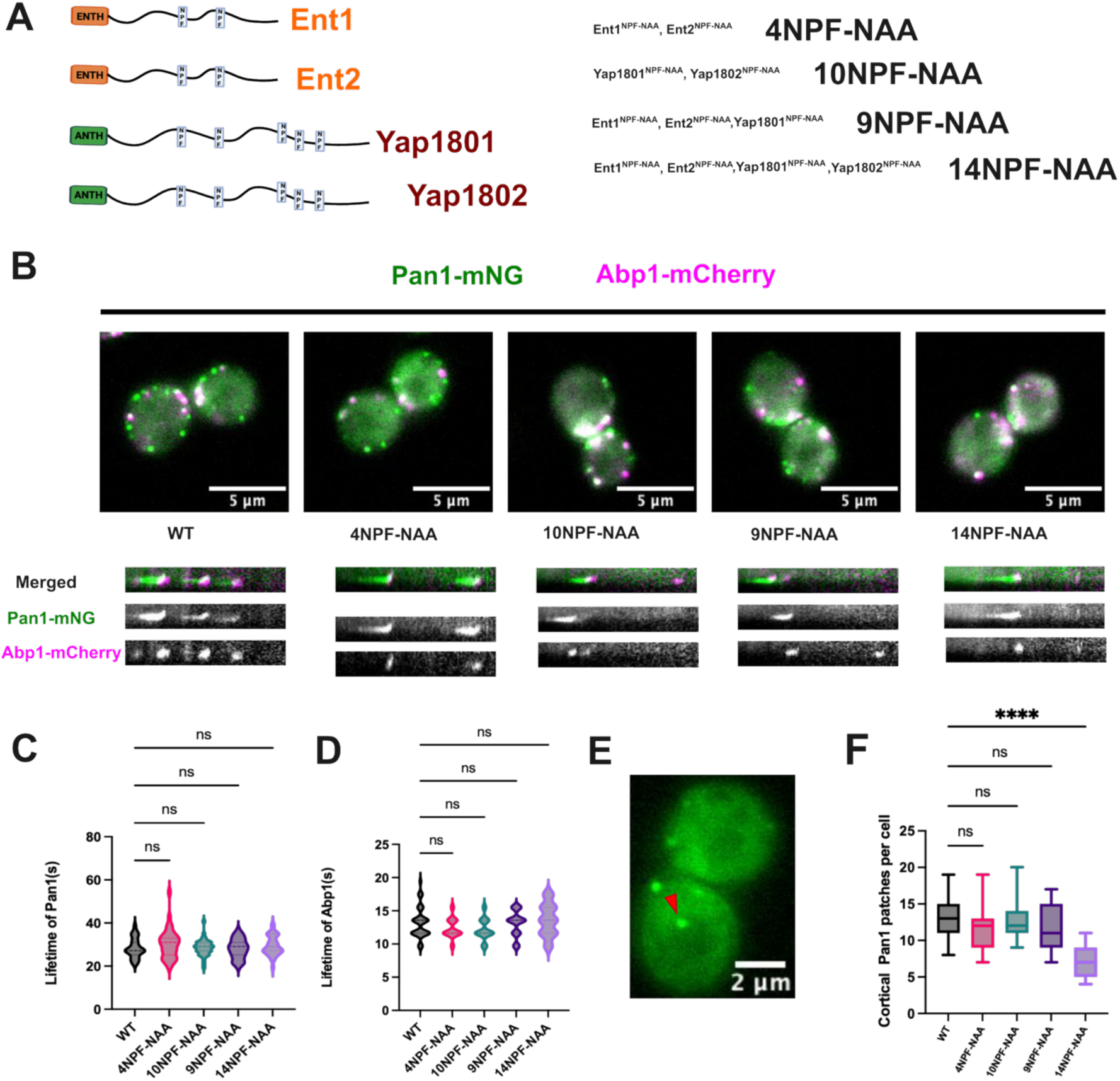
(**A**) Schematic representation of NPF motif-containing endocytic adaptors and mutant combinations generated in this study. NPF motifs in Ent1/2 and Yap1801/2 were mutated to NAA to disrupt EH domain interactions. Combined mutant strains containing 4, 9, 10, or 14 NPF-to-NAA substitutions are indicated. (**B**) Representative fluorescence images and corresponding kymographs of Pan1-mNG and Abp1-mCherry cortical patches in WT and indicated adaptor mutant background. Scale bars, 5 µm. **(C)** Quantification of Pan1 cortical patch lifetimes in WT and indicated adaptor mutant strains (one-way ANOVA, ns, P = 0.3241). n = 35 patches. **(D)** Quantification of Abp1-mCherry patch lifetimes in WT and indicated adaptor mutants. one-way ANOVA, ns). n = 35 patches. **(E)** Representative image showing Pan1 cytoplasmic assembly in the 14NPF-NAA mutant. **(F)** Quantification of cortical Pan1 puncta in wild-type and adaptor NPF mutant strains. (one-way ANOVA, ****P < 0.0001). n = 35 cells.

We next asked whether progressively disrupting the EH-NPF interactions altered Pan1 dynamics at endocytic sites. Surprisingly, mutation of all four NPF motifs across Ent1 and Ent2 (4NPF-NAA) had no detectable effect on Pan1 and Abp1 cortical patch dynamics compared to wild-type cells. Likewise, mutation of nine or ten NPF motifs in the combined adaptor backgrounds (9NPF-NAA and 10NPF-NAA) did not measurably alter Pan1 lifetime (Fig. 6A, B). Strikingly, even in the 14NPF-NAA strain, in which all fourteen NPF motifs across the Ent and Yap adaptor families were disrupted, Pan1 patch lifetimes remained indistinguishable from wild type (Fig. 6C). Thus, progressively eliminating the EH-NPF interactions failed to produce a corresponding progressive defect in Pan1 dynamics.

Pan1 lifetime analysis likewise showed normal lifetime of Pan1 patches (Fig. 6B). Consistent with this, Abp1-mCherry patch lifetimes remained unchanged across all NPF mutant backgrounds (Fig. 6D), indicating that disruption of EH-NPF interactions does not measurably alter progression of either the late coat or actin modules during endocytosis.

Despite the preservation of endocytic protein lifetime, mutation of all fourteen NPF motifs of adaptor proteins produced a striking reduction in the number of cortical Pan1 endocytic sites (Figure 6F). In these cells, Pan1 frequently formed cytosolic condensates rather than remaining restricted to cortical puncta (Figure 6E, Movie S3), resembling the condensates observed upon Pan1 overexpression. This is particularly interesting because partial disruption of adaptor NPF motifs produced no detectable effect on the number of endocytic sites or Pan1 localization. These observations suggest that adaptor mediated EH-NPF interactions redundantly determine Pan1 localization until a critical level of recruitment is lost, at which point Pan1 assembly becomes spatially deregulated, indicating that multivalent adaptor interactions contribute to spatial seeding of Pan1 at endocytic sites.

Because Ede1 also has EH domains that can interact with the NPFs and Ede1 deletion has been shown to cause reduction in late coat endocytic patches (Lu & Drubin, 2017) similar to what we observed in the 14NPF-NAA mutant, we next examined whether Ede1 localization was altered in the 14NPF-NAA background. Ede1 was recruited to cortical endocytic sites in 14NPF-NAA cells, indicating that endocytic-site localization of Ede1 was not abolished in these cells. However, in addition to these cortical puncta, Ede1 formed prominent cytoplasmic condensates (Fig. S5A). Thus, the reduction in cortical Pan1 sites in the 14NPF-NAA strain is unlikely to result simply from a loss of Ede1 from endocytic sites. Instead, disruption of adaptor-mediated EH-NPF interactions likely alters the recruitment of Pan1 directly, thereby also promoting its accumulation in ectopic condensates.

Together, these findings support a model in which the EH-NPF interactions function primarily as trigger for endocytic assembly rather than kinetic regulators of endocytosis. Rather than controlling endocytic progression, these interactions locally seed Pan1 assembly at cortical endocytic sites, thereby spatially restricting its self-assembly potential and ensuring productive assembly of the late endocytic coat.

## Discussion

The endocytic machinery assembles through a dense network of transient and multivalent interactions, yet how these interactions are organized into a spatially restricted and functional membrane-remodeling structure remains incompletely understood. Here, we show that the essential late endocytic scaffold Pan1 possesses an intrinsic capacity for higher-order self-assembly and that this capacity is subject to multiple levels of regulation. Our results support a model in which adaptor interactions spatially constrain Pan1 assembly to cortical endocytic sites, while a distal C-terminal region limits the extent of self-assembly. Together with the known phosphorylation-dependent regulation of Pan1, these findings suggest that Pan1 assembly is controlled not only in space but potentially also in its material state during endocytic progression.

Our findings extend a growing body of work implicating multivalent condensation in organizing the endocytic machinery. In budding yeast, condensation of the early scaffold Ede1 has been proposed to contribute to endocytic-site initiation, whereas reconstitution of the mammalian initiators Eps15 and Fcho1/2 demonstrated that weak, multivalent interactions can generate dynamic assemblies that concentrate downstream endocytic components (Day et al., 2021; Kozak & Kaksonen, 2022). Our results reveal a related organizational capacity in Pan1, which acts later in the endocytic pathway. Pan1 self-assembly depends on several regions distributed throughout the protein, including intrinsically disordered regions, EH2, and the central coiled-coil, suggesting that higher-order assembly emerges collectively from its multidomain architecture rather than from a single dedicated condensation module.

We interpret the assemblies produced by Pan1 overexpression as an experimental manifestation of this intrinsic self-assembly capacity rather than as direct representations of physiological endocytic patches. Several observations distinguish these structures from aggregates. They exhibit partial but rapid fluorescence recovery after photobleaching, undergo apparent fusion and fission, dissolve reversibly following treatment with 1,6-hexanediol or elevated temperature, and form membrane-free, ribosome-excluding structures at the ultrastructural level. These properties are consistent with a dynamic condensate-like assembly. However, Pan1 assembly does not fit with a simple equilibrium phase-separation model. Cytosolic Pan1 intensity increases with increasing promoter strength, indicating that condensate formation does not buffer the soluble Pan1 pool. One possible explanation is that Pan1 assembly depends on additional endocytic components, such as membrane lipids, clathrin-associated factors or actin regulators, which may become limiting upon overexpression. Under such conditions, part of the excess Pan1 could partition into condensates while the remaining protein accumulates diffusely in the cytoplasm. We have used the term condensation here to describe an intrinsic biochemical property of Pan1 revealed under elevated concentration rather than to conclude that physiological late endocytic coats are themselves biomolecular condensates.

Condensation and aggregation are not necessarily mutually exclusive states. Several intrinsically disordered and aggregation-prone proteins, including FUS, α-synuclein and tau, can form dynamic condensates that subsequently transition towards increasingly arrested or aggregated states (Patel et al., 2015; Ray et al., 2020; Wegmann et al., 2018). Pan1 may similarly occupy a broader self-assembly landscape. Consistent with this possibility, our sequence analysis identifies not only strong condensation-promoting features but also substantial aggregation-promoting propensity. Rather than representing contradictory predictions, these features may indicate that Pan1 has the potential to access molecular assemblies with different degrees of internal dynamics. This possibility is particularly intriguing given the changing physical requirements of the late endocytic coat. Early during late-coat assembly, rapid molecular exchange may facilitate recruitment and rearrangement of endocytic components. As the site matures and actin-dependent forces begin to deform the membrane a more constrained interaction network could provide increased structural and mechanical stability. We therefore speculate that Pan1 may shift along a continuum from a relatively dynamic, exchange-competent multivalent assembly towards a more constrained state as endocytosis progresses (Boinet et al., 2026). The known phosphorylation cycle of Pan1 provides a potential mechanism for regulating this behavior. Pan1 is phosphorylated by the endocytic kinases Ark1 and Prk1 during late stages of endocytosis, and preventing normal phosphorylation causes persistent endocytic structures, abnormal actin accumulations, and defective coat disassembly. Because many of these regulatory sites occur within interaction-rich regions of Pan1, phosphorylation could alter local charge, conformational behavior, or interaction strength, thereby destabilizing multivalent contacts that were advantageous during coat assembly. In this model, increasing molecular connectivity during coat maturation could permit Pan1 to enter a more constrained state, whereas late phosphorylation would promote escape from this state and facilitate network disassembly. The persistent structures observed when Ark1/Prk1-dependent regulation is impaired may therefore represent excessive stabilization of the Pan1-containing interaction network rather than a completely unrelated form of assembly (Sekiya-Kawasaki et al., 2003).

This interpretation suggests that the overexpression-induced Pan1 condensates and the abnormal structures produced by impaired Ark1/Prk1 signaling may be different assemblies. Aggregates are formed at endogenous expression level of Pan1 and contain clustered vesicles because of defective post-endocytic disassembly, whereas the Pan1 condensates described here are membrane-free structures induced by Pan1 overexpression. These phenotypes may illustrate different regions of a broader Pan1 assembly landscape in which multivalent interactions can generate structures with different compositions and degrees of molecular mobility. Testing this model will require direct measurements of endogenous Pan1 material properties, together with systematic manipulation of its phosphorylation state.

Our domain analysis provides further support for a distributed model of Pan1 assembly. Earlier structure-function analysis (Miliaras et al., 2004) showed that Pan1 can self-associate and suggested that its central coiled-coil contributes importantly to dimeric or oligomeric scaffold organization. Our results extend this model by showing that the coiled coil is necessary but not sufficient to explain Pan1 self-assembly. In addition to the coiled-coil, the N-terminal IDR, EH2, and a central disordered region are independently required for efficient condensate formation, indicating that Pan1 assembly emerges through cooperation between several regions distributed across the protein. Conversely, removal of a distal C-terminal region enhanced both condensate size and number. This region contains interaction sites for the SH3 domains of the type I myosins Myo3 and Myo5 (Barker et al., 2007), raising the possibility that interactions involving this region normally constrain Pan1 self-assembly. The enhanced condensation upon deletion of distant C-terminal region is interesting in light of the transition from coat maturation to actin-driven membrane invagination. The type I myosins are recruited as the endocytic machinery enters its force-generating phase, raising the possibility that interactions with the Pan1 C-terminal region reorganize or constrain the scaffold as the molecular requirements of the coat change. Day et al. (Day et al., 2021) showed that altering interaction strength within the Eps15/Fcho system can shift assemblies towards progressively more arrested states. Although the underlying mechanism may differ for Pan1, our results similarly suggest that modulation of individual interaction interfaces could tune the physical organization of a multivalent scaffold rather than simply determining whether assembly occurs.

Importantly, the same intrinsically disordered regions required for overexpression-induced Pan1 condensation also influence the dynamics of endogenous Pan1 and Sla1, linking the self-assembly assay to physiological late-coat organization. Deletion of these regions prolongs the lifetime of Pan1- and Sla1-containing endocytic sites without equivalently altering the dynamics of the actin marker Sac6, suggesting that distributed Pan1 interactions contribute specifically to organization and remodeling of the late coat. Thus, the condensation assay does not merely reveal an overexpression phenotype but identifies molecular regions whose interaction properties are also relevant at endogenous expression level.

Precise disruption of EH-NPF interactions reveals a second and mechanistically distinct level of Pan1 regulation: seeding of Pan1 assembly at the endocytic sites. Previous studies using adaptor- deletion backgrounds established important roles for the epsins and Yap180 proteins in Pan1 lifetime and endocytic progression (Maldonado-Báez et al., 2008). However, these adaptors contribute to membrane association, cargo recognition, clathrin recruitment and interactions with numerous other proteins in addition to their NPF-mediated contacts with Pan1. Removing entire adaptors therefore makes it difficult to isolate the specific contribution of the EH-NPF interaction network. By progressively mutating the canonical NPF motifs within full-length Ent1, Ent2, Yap1801 and Yap1802, we were able to disrupt the NPF-EH domain interaction network while retaining other adaptor functions. Surprisingly, removal of increasing numbers of NPF motifs produced no significant effect on the lifetime of Pan1 or Abp1. In the 14NPF-NAA background, however, the number of cortical Pan1 patches declined sharply and Pan1 accumulated in ectopic cytoplasmic assemblies. The principal function of the adaptor-Pan1 interaction network therefore appears to be less about determining the lifetime of Pan1 once a productive site has formed and more about ensuring the correct localization of Pan1 assembly.

The strongly nonlinear response to progressive NPF removal further reveals substantial redundancy within this system. Partial disruption can be buffered by the remaining interactions, whereas endocytic site seeding fails once adaptor engagement falls below a critical level. However, once Pan1 successfully assembles at a cortical site, much of the downstream endocytic process can still proceed normally.

One possible explanation for this robustness in Pan1 assembly is that canonical NPF motifs represent preferred but not exclusive ligands for EH domains. Recent NMR analysis of mammalian Eps15 showed that its EH domains can recognize several non-canonical phenylalanine-containing motifs in addition to NPF and can engage intrinsically disordered regions lacking canonical NPF sequences, generally with somewhat lower affinity (Papagiannoula et al., 2025). Although equivalent binding promiscuity has not yet been shown directly for Pan1, such interactions could help explain why Pan1 can still assemble and function normally at some endocytic sites. In this view, Pan1 recruitment could emerge from the collective action of canonical NPF contacts together with weaker non-canonical interactions, producing a highly redundant seeding system.

Together, our findings suggest that Pan1 self-assembly is regulated along at least three interconnected dimensions. First, distributed interactions within Pan1 provide an intrinsic capacity for higher-order assembly. Second, EH-NPF interactions with membrane-associated adaptors determine where this assembly is initiated, effectively seeding Pan1 at cortical endocytic sites. Third, interactions involving the C-terminal myosin-binding region, together with phosphorylation-dependent remodeling, may determine how extensively and how persistently the resulting network assembles. These regulatory layers would allow Pan1 to exploit the organizational advantages of multivalent self-assembly while preventing non-productive accumulation in the cytoplasm.

This model places Pan1 alongside Ede1 and the mammalian initiators Eps15 and Fcho1/2 as an endocytic scaffold whose function is intimately connected to regulated higher-order assembly, while extending this principle to a distinct stage of the pathway. Whereas condensation of Ede1 has been linked to endocytic-site initiation, Pan1 functions during late-coat maturation and the transition to actin-driven membrane invagination. The coexistence of dynamic condensation-promoting and aggregation-promoting features may therefore be especially relevant for Pan1, allowing its interaction network to remain rearrangeable during assembly while acquiring greater constraint as mechanical demands increase, before being reset through phosphorylation-dependent disassembly.

Our work supports a model in which spatially seeded and dynamically regulated multivalent self-assembly organizes the late endocytic coat. More broadly, it suggests that cellular scaffolds need not occupy a single fixed material state. Instead, their physical organization may be continuously tuned by localization signals, interaction partners, and post-translational modifications, allowing dynamic molecular networks to acquire and subsequently release the structural stability required for cellular function.

## Materials and Methods

### Yeast strains, growth conditions and strain construction

Yeast strains and plasmids used in this study are listed in Table S1. All strains were derived from the parental *Saccharomyces cerevisiae* strain DDY1102 (*his3-Δ200, leu2-3,112, ura3-52, lys2-801*). Cells were grown in rich medium at 24 °C. Gene deletions, fluorescent protein tagging and introduction of mutant alleles were performed by homologous recombination using PCR-amplified cassettes as described by (Janke et al., 2004).

Pan1 truncation mutants were generated in a diploid background because *PAN1* is essential. One copy of *PAN1* was first deleted from the homozygous diploid strain and replaced with a LEU2 selection cassette, generating a heterozygous *PAN1/pan1Δ::LEU2* strain. The deletion allele was subsequently replaced with the corresponding truncated *PAN1* allele linked to a natNT2 cassette. Transformants were selected on medium containing nourseothricin. Correct integration of each truncation construct was initially verified by PCR amplification of the modified genomic locus. The amplified region was subsequently analyzed by Oxford Nanopore Technologies sequencing to confirm the identity and integrity of the introduced truncation.

### Plasmid construction

Plasmids used in this study are listed in Table S2. DNA fragments were amplified by PCR using the appropriate primers and assembled into the corresponding plasmid backbones using HiFi DNA assembly. The resulting constructs were transformed into *Escherichia coli*, purified and verified by DNA sequencing before transformation into yeast.

### Live-cell fluorescence microscopy

Yeast cells were grown at 24 °C to an OD of 0.3-0.8 in low-fluorescence synthetic dropout medium lacking tryptophan. Cells were immobilized on coverslips coated with 1 mg mL −1 concanavalin A.

### Widefield microscopy

Widefield fluorescence images were acquired using an Olympus IX81 microscope equipped with a 100×/1.45 NA objective, an ORCA-ER CCD camera (Hamamatsu) and an X-CITE 120 PC metal-halide illumination source (EXFO). EGFP- and mCherry-tagged proteins were imaged using U-MGFPHQ and U-MRFPHQ filter sets, respectively. Image acquisition was controlled using Micro-Manager software.

### Spinning-disk confocal microscopy

Spinning-disk confocal imaging for Figure 1 was performed at the Photonic Bioimaging Center, University of Geneva, using a Nikon Eclipse Ti1 microscope equipped with a Yokogawa CSU-W1 spinning-disk unit, a 100×/1.49 NA objective and a Prime 95B sCMOS camera (Photometrics). eGFP signal was excited using 488 nm laser.

### Fluorescence recovery after photobleaching

FRAP experiments were performed on Pan1 condensates using an Olympus IX83 microscope equipped with an iLas2 targeted illumination system. Pan1 condensates were photobleached using a 488 nm laser. Fluorescence recovery within the bleached region was further monitored by time-lapse imaging under low-intensity illumination. Recovery curves were normalized to the pre-bleach fluorescence intensity and corrected for background fluorescence and photobleaching.

### Correlative light and electron microscopy

Correlative light and electron microscopy was performed as described previously (Kukulski et al., 2011). *Saccharomyces cerevisiae* cells expressing Pan1 tagged with yeGFP and expressed under the TEF promoter were grown in low-fluorescence medium at 24 °C to logarithmic phase. Cells were collected by vacuum filtration and cryo-fixed by high-pressure freezing using a Leica EM ICE high-pressure freezer.

Freeze substitution was performed using a Leica AFS2 system. Samples were incubated in glass-distilled acetone containing 0.1% (w/v) uranyl acetate first on dry ice with shaking for 2-3 hours and then at −90 °C for 16 h. The temperature was subsequently increased to −45 °C at a rate of 5 °C h⁻¹. Samples were washed with acetone and infiltrated with increasing concentrations of Lowicryl resin in acetone (10%, 25%, 50% and 75%; 4 h each) while the temperature was gradually increased to −25 °C. Samples were then incubated in 100% Lowicryl resin, which was exchanged three times at 10 h intervals, and polymerized under ultraviolet light at −25 °C for 48 h. The temperature was subsequently increased to 20 °C at 5 °C h⁻¹, and ultraviolet light polymerization was continued for an additional 48 h. Sections of around 300 nm thickness were cut using an ultramicrotome equipped with a diamond knife and collected on carbon-coated 200-mesh copper grids. Fluorescence imaging of resin sections and subsequent electron tomography followed by correlation were performed according to (Ivanović et al., 2026).

### Image analysis

Protein lifetimes were measured from time-lapse movies acquired by widefield fluorescence microscopy. Movies were corrected for photobleaching and background signal using in-lab Fiji plugin. Fluorescence intensity profile was extracted for individual endocytic events. The start and end of each event was defined as the point at which the fluorescence signal was above and then returned to the baseline intensity measured in regions without detectable endocytic events. Protein lifetime was calculated as the interval between these two time points.

The number of endocytic sites was quantified in fully budded cells. Endocytic sites were counted manually from a single image acquired at the equatorial plane of each cell.

Cytoplasmic fluorescence intensity was quantified manually in ImageJ using 5 × 5-pixel square regions of interest positioned away from condensates and vacuoles.

### Western blotting

Protein extracts were prepared from 5 mL exponentially growing yeast cultures by adding 300 µL ice-cold trichloroacetic acid. Cells were collected by centrifugation, washed with cold acetone and dried in a vacuum concentrator. Cell pellets were resuspended in 100 µL urea buffer containing 25 mM Tris-HCl, pH 6.8, 6 M urea and 1% SDS, and disrupted by vigorous shaking with 200 µL glass beads. Samples were heated at 95 °C for 5 min, mixed with 100 µL 2× SDS sample buffer and centrifuged at 16,000 × *g* for 5 min.

Proteins were separated on 4-20% precast polyacrylamide gels (Bio-Rad) and transferred onto nitrocellulose membranes using an iBlot 2 system (Thermo Fisher Scientific). Membranes were blocked for 30 min in 5% skimmed milk prepared in TBS-T and incubated overnight at 4 °C with primary antibodies for GFP (abcam, ab291) and PGK1(abcam, ab113687) in 1:2000 dilution. After washing with TBS-T, membranes were incubated for 1 h at room temperature with HRP-conjugated secondary antibody (abcam, ab6728-1) in 1:5000 dilution. Protein bands were detected using an enhanced chemiluminescence substrate and imaged by chemiluminescence detector.

### Phalloidin staining

Cells were grown overnight to mid-logarithmic phase fixed by adding paraformaldehyde to a final concentration of 3.7% and incubating for around 60 min at 30°C with shaking. Fixed cells were collected by centrifugation at and washed three times with PBS. Cells were then washed and resuspended in PBT.

Filamentous actin was stained with Alexa Fluor 568-conjugated phalloidin at a final concentration of 2.5 µM for 1 h at room temperature in the dark. Cells were then washed and resuspended in PBS and mounted on concanavalin A coated coverslips.

### FM4-64 internalisation assay

Cells were grown to mid-logarithmic phase in synthetic complete medium lacking tryptophan, prepared for live-cell imaging as described above and mounted on concanavalin A coated coverslips. After the sample was positioned on the IX81 microscope, the medium was replaced with SC-Trp containing 5 µM FM4-64. Time-lapse imaging was initiated immediately after addition of the dye, and FM4-64 internalization was followed at the indicated time points.

### Yeast two hybrid assay

Yeast-Two Hybrid assay was performed by cloning gene fragments of interest into Sal1 digested pMM5 and pMM6 plasmids, which contain the LexA DNA binding domain and Gal4 Activation domain respectively. The resulting plasmids were transformed into *Saccharomyces cerevisiae* strains SGY37 (MATa) and YPH500 (MATα).

To map the regions of Pan1 involved in self-interaction, different Pan1 fragments were cloned into Y2H vectors and tested for their ability to interact with each other. Yeast strains carrying the respective constructs were mated on YPD plates and incubated at 30°C for 2 days. The resulting diploids were then replica-plated onto SC-His-Leu double-selection plates and incubated at 30°C for an additional 2 days. Protein-protein interactions were subsequently assessed by measuring β-galactosidase activity.

For detection of β-galactosidase activity, the plates were overlaid with freshly prepared X-Gal solution containing 500 mM sodium phosphate buffer (pH 7.0), 10% SDS, 1 M KCl, 1 M MgCl₂, 0.04% X-Gal, and 0.4% low-melting-point agarose. Interaction between LexA-DBD-Pan1 and Gal4-AD-Pan1 fragments reconstitutes the transcriptional activation system, resulting in expression of the *lacZ* reporter gene encoding β-galactosidase. β-galactosidase activity converts X-Gal into a blue-coloured product, providing a visual readout of the interaction. Plates were scanned and documented after 10-12 h of incubation with the X-Gal overlay mixture.

### Statistical analysis

Statistical analysis was performed using Graph pad prism software.

### Yeast strains and plasmids

All the yeast strains and plasmids used in the study are listed in Table S1 and S2 in supporting information file.

## Acknowledgments

We thank all the members of the Kaksonen lab for their feedback, Christopher Toret and Anne Laure Boinet for reading the manuscript, and Afia Mougamadoubougary for her assistance. We thank the DCI Geneva (CryoGEnic) facility at the University of Geneva as well as the EM facility at the Institute of Anatomy and the Microscopy Imaging Center of the University of Bern for access to equipment and technical support. We also thank Dr. Christoph Bauer and Dr. Andrew Howe for assistance with EM sample preparation and data collection. We are grateful to Prof. Gislene Pereira for generously providing the Y2H strains and to Prof. David Drubin for sharing the *ark1Δ prk1-as3* strain used in the study. Work in the Kukulski lab was supported by the SNSF NCCR TransCure 185544 and SNSF project grant 201158 and in the Kaksonen lab by the SNSF project grant 212288.

