## Supporting information for "Adaptor interactions trigger Pan1 self-assembly during clathrin-mediated endocytosis in budding yeast"

Figures

Fig. S1.

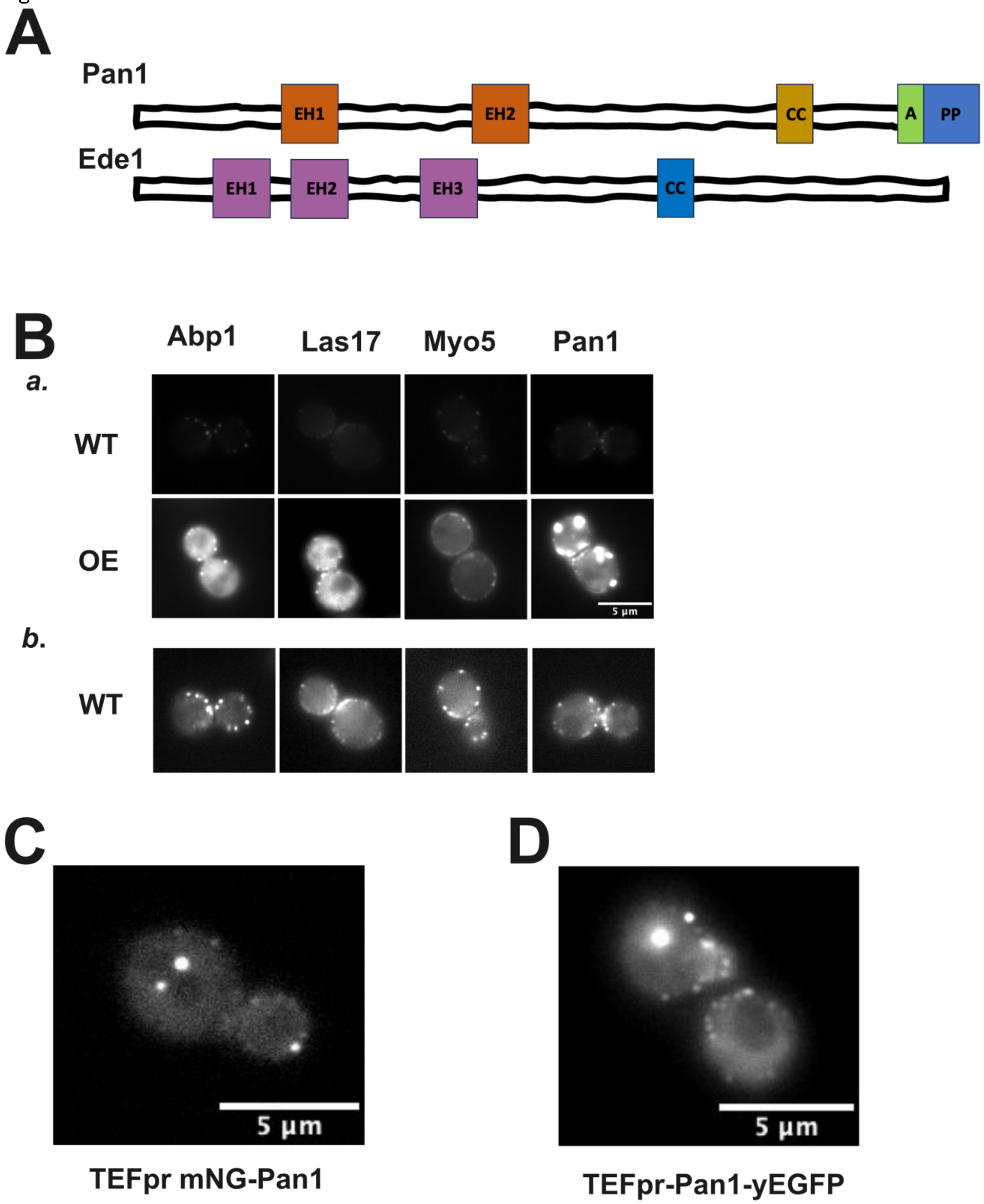


(A) Domain organisation of Pan1 and Ede1. (B) a. Representative fluorescence images comparing the localization of endocytic proteins in wild type cells (WT) and upon over-expression (OE). b. Images are shown with independently adjusted brightness and contrast to visualize the comparatively dim endocytic sites. (C) Fluorescence image of mNG-tagged Pan1 expressed from the TEF1 promoter. (D)

Representative fluorescence image of C-terminally tagged Pan1-yEGFP expressed from the TEF1 promoter.

Fig. S2.

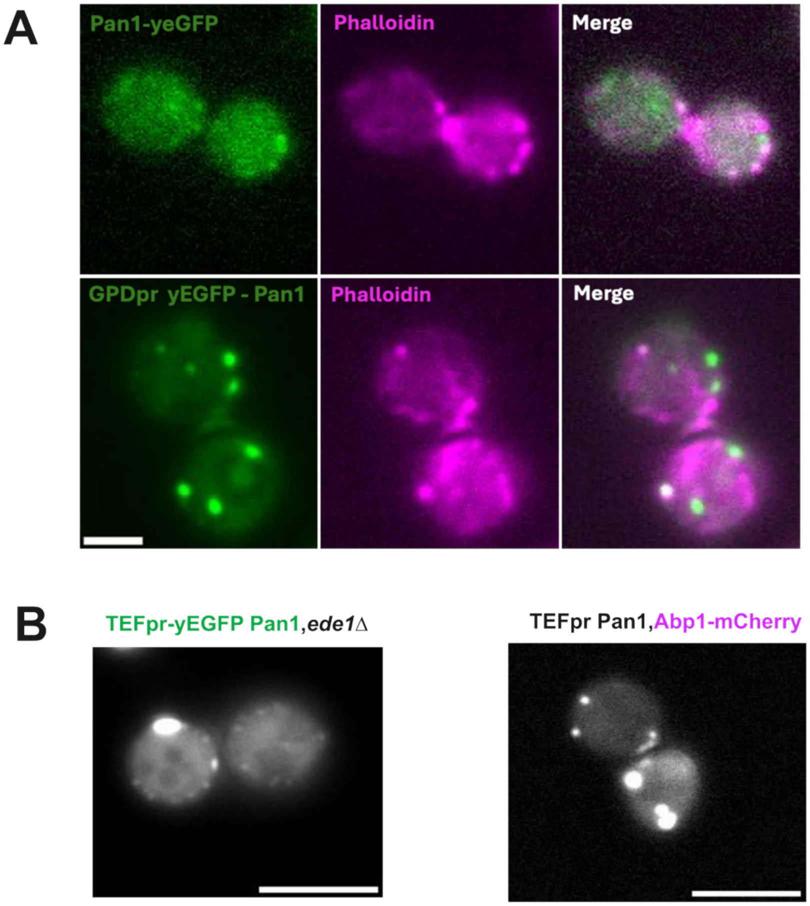


**(A)** Representative fluorescence images of Pan1 condensates stained with phalloidin. (B) Representative fluorescence image of TEFpr yEGFP-Pan1 in an ede1Δ background. (C) Representative fluorescence image of Abp1-mCherry in cells overexpressing untagged Pan1. Scale bars, 5 μm.

Fig. S3.

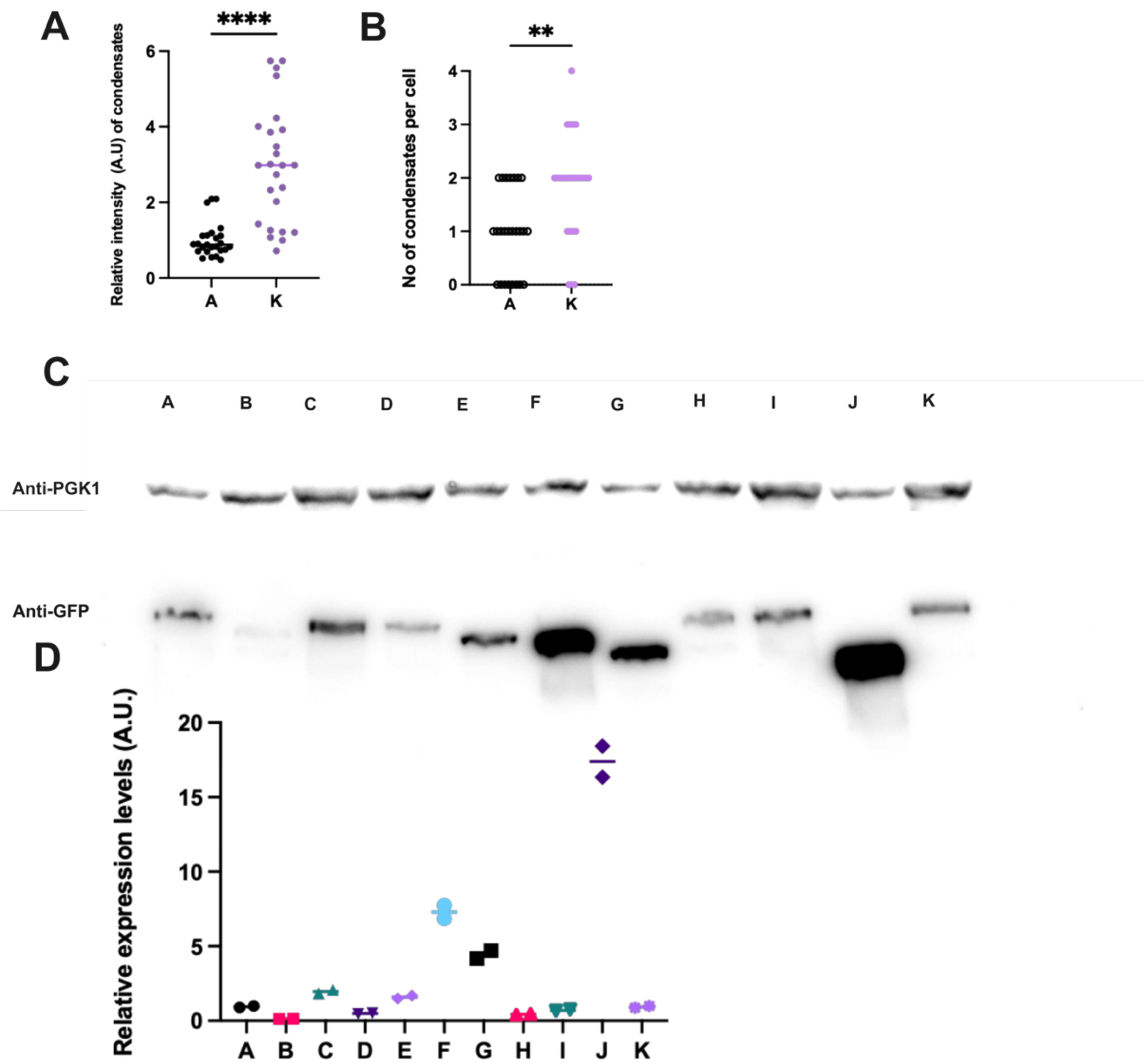


A) Quantification of the relative fluorescence intensity of Pan1 condensates in the indicated strains discussed in Fig. 4. The condensation-enhancing mutant K (PAN1 /TEFpr-yEGFP pan1(1310-1480Δ)) exhibited significantly higher condensate fluorescence intensity than A (TEFpr-yEGFP PAN1 / pan1Δ) (two-tailed unpaired t test, $P < 0.0001$; n = 25). (B) Quantification of the number of Pan1 condensates per cell in the indicated strains. Mutant K formed significantly more condensates than mutant A (two-tailed unpaired t test, $P = 0.0011$; n = 25 cells). (C) Representative immunoblot showing expression of GFP-tagged full-length Pan1 and the truncation mutants analysed in Fig. 4. GFP-tagged Pan1 constructs were detected using an anti-GFP antibody. PGK1 served as a loading control. (D) Quantification of GFP signal normalized to PGK1 for each Pan1 construct shown in (C).

Fig. S4.

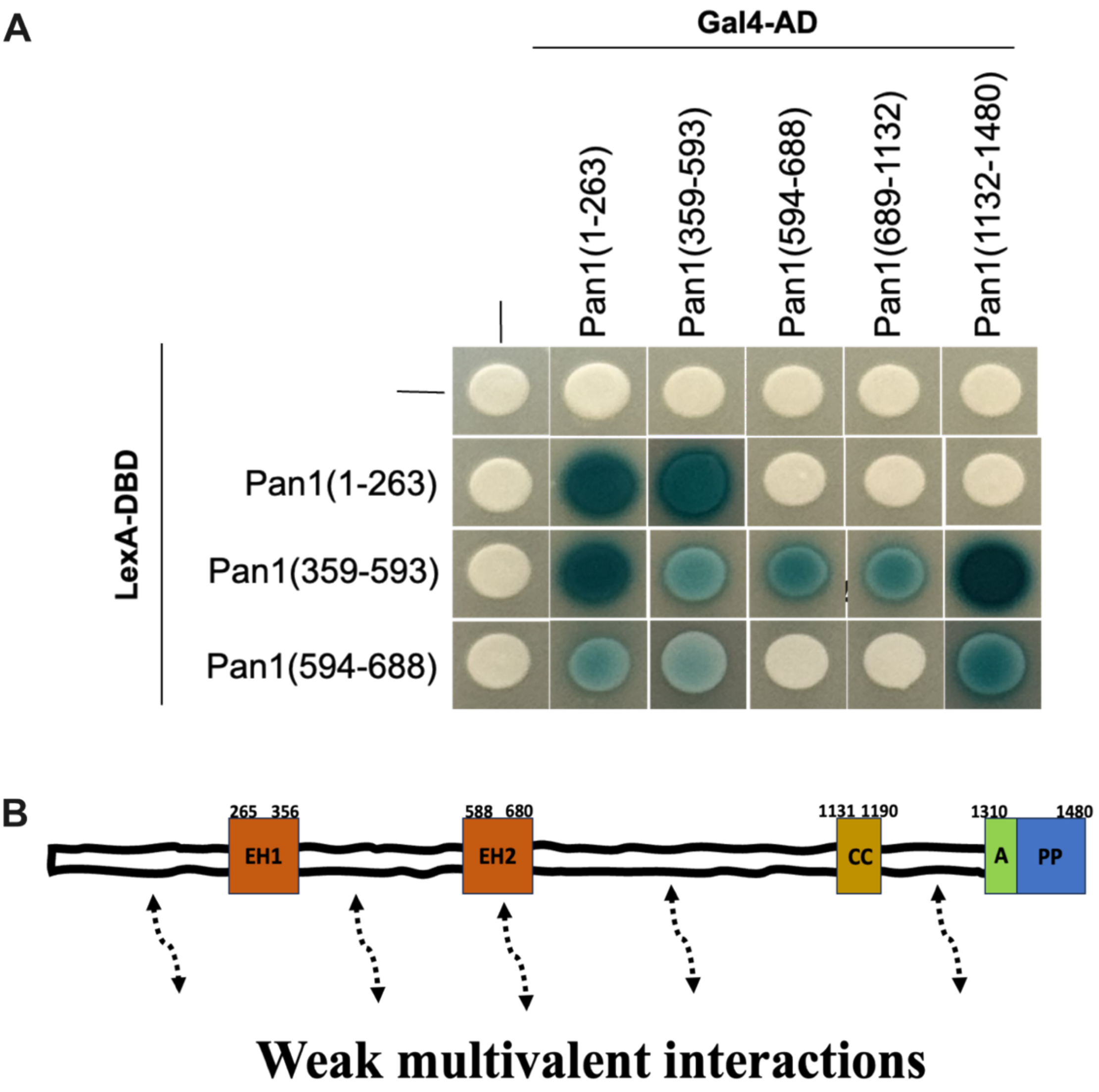


(A) Yeast two-hybrid analysis of interactions between distinct Pan1 fragments. Growth and reporter activation indicate multiple interaction interfaces distributed across the Pan1 sequence, consistent with weak multivalent self-association. (B) Schematic model illustrating cooperative multivalent interactions within Pan1 sequence.

Fig. S5.

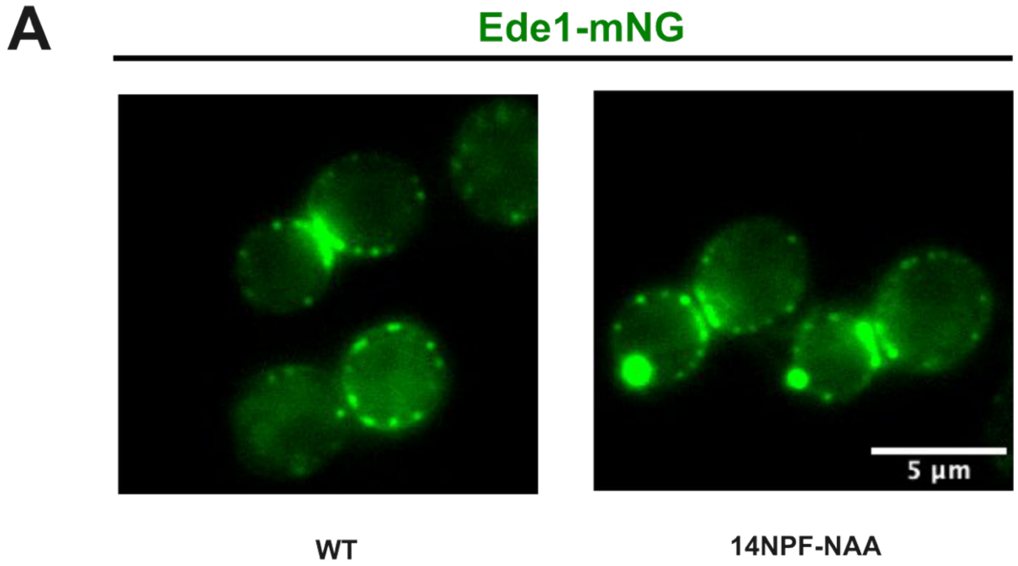


(A) Representative fluorescence images of Ede1-mNG localization in wild-type and 14NPF-NAA cells. Scale bars, 5 µm.

Tables

Table S1. Saccharomyces cerevisiae strains used in the study

| | |
|---|---|
| MKY0676 | MATa, his3200, leu2-3,112, ura3-52, lys2-801, Pan1-EGFP::HIS3MX6 |
| MKY4705 | MATa, his3200, leu2-3,112, ura3-52, lys2-801, GPDpr-yeGFP-Pan1::natNT2 |
| MKY4704 | MATa, his3200, leu2-3,112, ura3-52, lys2-801, TEFpr-yeGFP-Pan1 ::natNT2 |
| MKY4703 | MATa, his3200, leu2-3,112, ura3-52, lys2-801, CYCpr-yeGFP-Pan1::natNT2 |
| MKY4865 | MATa, his3200, leu2-3,112, ura3-52, lys2-801, GPDpr-yEGFP-PAN1::natNT2, Sac6-mCherry::KANMX4 |
| MKY4867 | MATa, his3200, leu2-3,112, ura3-52, lys2-801, GPDPr- yEGFP-Pan1::natNT2, Abp1-mCherry::KANMX4 |
| MKY4869 | MATa, his3200, leu2-3,112, ura3-52, lys2-801, GPDPr- yEGFP-Pan1::natNT2, Ede1-mCherry::KANMX4 |
| MKY4886 | MATa, his3200, leu2-3,112, ura3-52, lys2-801,GPDPr- yEGFP-Pan1 ::natNT2, Sla1-mCherry::KANMX4 |
| MKY4971 | MATa, his3200, leu2-3,112, ura3-52, lys2-801, GPDPr- yEGFP-Pan1::natNT2, Myo5-mCherry::KanMX4 |
| MKY5572 | MAT?, his3200, leu2-3,112, ura3-52, lys2-801, GPDpr-yeGFP-PAN1::natNT2, Yap1801-mCherry::KanMX4 |
| MKY5468 | MATalpha, his3200, leu2-3,112, ura3-52, lys2-801, GPDpr-yeGFP-Pan1::natNT2, Rvs167-mCherry::KANMX4 |
| MKY5566 | MATa, his3200, leu2-3,112, ura3-52, lys2-801, GPDpr-yeGFP-Pan1::natNT2, SLA2-mCherry::kanMX4 |
| MKY5568 | MATalpha, his3200, leu2-3,112, ura3-52, lys2-801, GPDpr-yeGFP-Pan1::natNT2, Syp-1-mCherry::kanMX4 |
| MKY5570 | MATa, his3200, leu2-3,112, ura3-52, lys2-801, GPDpr-yeGFP-Pan1::natNT2, Ent1-mCherry::kanMX4 |
| MKY5587 | MATa, his3200, leu2-3,112, ura3-52, lys2-801, GPDpr-yeGFP-Pan1::natNT2, Ark1-mCherry::kanMX4 |
| MKY4986 | his3-Δ200, leu2-3,112, ura3-52, ADE2, lys2-801(oc), ark1Δ::cgHIS3, prk1-as3::URA3 |
| MKY5724 | MATa/alpha, his3200, leu2-3,112, ura3-52, lys2-801, TEFpr-yeGFP-Pan1 ::natNT2 |
| MKY5795 | MATa/alpha, his3200, leu2-3,112, ura3-52, lys2-801, TEFpr-yeGFP-Pan1(2-265Δ) ::natNT2 |
| MKY5796 | MATa/alpha, his3200, leu2-3,112, ura3-52, lys2-801, TEFpr-yeGFP-Pan1(265-356Δ) ::natNT2 |
| MKY5721 | MATa/alpha, his3200, leu2-3,112, ura3-52, lys2-801, TEFpr-yeGFP-Pan1(588-680Δ) ::natNT2 |
| MKY5722 | MATa/alpha, his3200, leu2-3,112, ura3-52, lys2-801, TEFpr-yeGFP-Pan1(265-680Δ) ::natNT2 |
| MKY5723 | MATa/alpha, his3200, leu2-3,112, ura3-52, lys2-801, TEFpr-yeGFP-Pan1(680-1130Δ) ::natNT2 |
| MKY5794 | MATa/alpha, his3200, leu2-3,112, ura3-52, lys2-801, TEFpr-yeGFP-Pan1(2-680Δ) ::natNT2 |
| MKY5730 | MATa/alpha, his3200, leu2-3,112, ura3-52, lys2-801, TEFpr-yeGFP-Pan1(1191-1310Δ) ::natNT2 |
| MKY5797 | MATa/alpha, his3200, leu2-3,112, ura3-52, lys2-801, TEFpr-yeGFP-Pan1(1130-1190Δ) ::natNT2 |
| MKY5798 | MATa/alpha, his3200, leu2-3,112, ura3-52, lys2-801, TEFpr-yeGFP-Pan1(792-1480Δ) ::natNT2 |
| MKY5808 | MATa/alpha, his3200, leu2-3,112, ura3-52, lys2-801, TEFpr-yeGFP-Pan1(1310-1480Δ) ::natNT2 |

| | |
|---|---|
| MKY6012 | MATα, his3-Δ200, leu2-3,112, ura3-52, lys2-801, pan1(Δ1-263)::ClonNat, Sla1-EGFP::HIS3MX6, Sac6-mCherry::KANMX4 |
| MKY6006 | MATα, his3-Δ200, leu2-3,112, ura3-52, lys2-801, pan1-(Δ1-263)-EGFP::ClonNat |
| MKY6008 | MATa, his3-Δ200, leu2-3,112, ura3-52, lys2-801, pan1-(Δ356-588)-EGFP::ClonNat |
| MKY5979 | MATα, his3-Δ200, leu2-3,112, ura3-52, lys2-801, pan1-(Δ356-588)-EGFP::ClonNat, Sac6-mCherry::KANMX4 |
| MKY6003 | MATα, his3-Δ200, leu2-3,112, ura3-52, lys2-801, pan1-(Δ356-588)-EGFP::ClonNat, Sla1-mCherry::KANMX4 |
| MKY5143 | MAT?, his3-Δ200, leu2-3,112, ura3-52, lys2-801, Pan1-mNG::HIS3MX6, ABP1-mCherry::KANMX4 |
| MKY5504 | MAT?, his3200, leu2-3, 112, ura3-52, lys2-801, ENT1-NAA::URA3, Ent2-NAA::hpH, ABP1-mCherry::kanMX4, Pan1-mNeonGreen::HIS3MX6 |
| MKY5503 | MAT?, his3200, leu2-3, 112, ura3-52, lys2-801, Yap1801-NAA::NatNT2, Yap1802-NAA::LYS2, ABP1-mCherry::kanMX4, Pan1-mNeonGreen::HIS3MX6 |
| MKY5500 | MAT?, his3200, leu2-3, 112, ura3-52, lys2-801, ENT1-NAA::URA3, Ent2-NAA::hpH, Yap1801-NAA::NatNT2, ABP1-mCherry::kanMX4, Pan1-mNeonGreen::HIS3MX6 |
| MKY5968 | MATalpha, his3200, leu2-3, 112, ura3-52, lys2-801, ENT1-NAA::URA3, Ent2-NAA::hpH, Yap1801-NAA::NatNT2, Yap1802-NAA::LYS2, ABP1-mCherry::kanMX4, Ede1-mNeonGreen::HIS3MX6 |
| MKY5499 | MAT?, his3200, leu2-3, 112, ura3-52, lys2-801, ENT1-NAA::URA3, Ent2-NAA::hpH, Yap1801-NAA::NatNT2, Yap1802-NAA::LYS2, ABP1-mCherry::kanMX4, Pan1-mNeonGreen::HIS3MX6 |
| MKY5114 | MATa, his3-Δ200, leu2-3,112, ura3-52, lys2-801, Ede1-mNeonGreen::HIS3MX6 |
| MKY5973 | MATalpha, his3-Δ200, leu2-3,112, ura3-52, lys2-801, TEFpr-Pan1::Nat, ABP1-mCherry::kanMX |
| MKY4936 | MATa, his3200, leu2-3,112, ura3-52, lys2-801,Pan1-EGFP::HIS3MX6, GPD pr:: natNT2 |
| MKY5975 | MATalpha, his3-Δ200, leu2-3,112, ura3-52, lys2-801, TEFpr-mNG-Pan1::Nat |
| MKY4824 | MATa, his3200, leu2-3,112, ura3-52, lys2-801, TEFpr-yeGFP-Pan1::natNT2, EDE1delta::NAT |
| MKY5964 | MATα, his3-Δ200, leu2-3,112, ura3-52, lys2-801, TEFpr-Pan1::Nat |
| MKY0106 | MATa, his3Δ-200, leu2-3,112, ura3-52, lys2-801, SLA1-mCherry::kanMX4 |
| MKY3996 | MATα, his3-Δ200, leu2-3,112, ura3-52, lys2-801, SAC6-mCherry::KanMX4 |
| MKY5576 | MATa leu2 ADE2 ura3-52::URA3-lexA-op-LacZ his3 trp1 |
| MKY5577 | MATα ura3-52 lys2-801amber ade2-101ochre trp1Δ63 his3Δ200 leu2Δ1 |

Table S2. Plasmids used in the study

| | |
|---|---|
| pMK0045 | pYM-N17 |
| pMK0010 | pYM-N21 |
| pMK0508 | pFA6a-Leu2 |
| pMK0553 | pYM-N21-Pan1 |
| pMk0558 | pYM-N21-Pan1 2-680 Δ |
| pMk0559 | pYM-N21-Pan1 2-265 Δ |
| pMk0560 | pYM-N21-Pan1 265-356 Δ |
| pMk0561 | pYM-N21-Pan1 588-680 Δ |
| pMk0562 | pYM-N21-Pan1 265-680 Δ |
| pMk0563 | pYM-N21-Pan1 1131-1190 Δ |
| pMk0564 | pYM-N21-Pan1 680-1480 Δ |
| pMk0565 | pYM-N21-Pan1 589-1244 Δ |
| pMk0566 | pYM-N21-Pan1 1191-1310 Δ |
| pMk0567 | pYM-N21-Pan1 681-1130 Δ |
| pMk0568 | pYM-N21-Pan1 1311-1480 Δ |
| pMk0569 | pYM-N21-Pan1 792-1480 Δ |
| pMK0583 | pYM-N21-Pan1 356-588 Δ |
| pMK0699 | pYM-N21 (TEF-Promotor)-mNG |
| pMK0471 | Ent1 NAA_pUC57 (Genescript) |
| pMK0472 | Ent2 NAA_pUC57 (Genescript) |
| pMK0473 | Yap1801NAA_pUC57(Genescript) |
| pMK0474 | Yap1802NAA_pUC57(Genescript) |
| pMK0019 | pFA6a-natNT2 |
| pMK0003 | pFA6a-EGFP-His3MX6 |
| pMK0333 | pFA6a-mNG-His3MX6 |
| pMK0695 | pFA6a-Pan1-IDR2Δ-EGFP-natNT2 |
| pMK0694 | pFA6a-Pan1-IDR1Δ-EGFP-natNT2 |
| pMK0005 | pFA6a-mCherry-KanMX4 |
| pMK0023 | pYM-N11 |
| pMK0732 | pMM5-Pan1(1-263) |
| pMK0733 | pMM5-Pan1(264-358) |
| pMK0734 | pMM5-Pan1(359-593) |
| pMK0735 | pMM5-Pan1(594-688) |
| pMK0736 | pMM5-Pan1(689-1131) |
| pMK0737 | pMM5-Pan1(1132-1480) |
| pMK0738 | pMM6-Pan1(1-263) |
| pMK0739 | pMM6-Pan1(264-358) |
| pMK0740 | pMM6-Pan1(359-593) |
| pMK0741 | pMM6-Pan1(594-688) |
| pMK0742 | pMM6-Pan1(689-1131) |
| pMK0743 | pMM6-Pan1(1132-1480) |

Movies (separate files).

Movie S1: One hour movie of TEFpr-yEGFP Pan1.

Movie S2: Tomogram for Pan1 condensates.

Movie S3: Movie showing Pan1 assemblies in cytoplasm at endogenous level in 14NPF-NAA mutant.
